# Yes, they’re all individuals: Hierarchical models from repeat measurement data improve estimates of tree growth and size

**DOI:** 10.1101/2024.08.27.609830

**Authors:** Theresa O’Brien, David Warton, Daniel Falster

**Affiliations:** University of New South Wales

**Keywords:** Bayesian models, repeat measurement surveys, forest ecology, growth modeling

## Abstract

1. Repeat measurement surveys of tree size are used in forests to estimate growth behaviour, biomass, and population dynamics. Although size is measured with error, and individuals vary in their growth trajectories, current size-based growth modelling approaches do not usually or fully account for both of these features, and therefore under-utilise available data.
2. We present a new method that leverages the auto-correlation structure of repeat surveys into a hierarchical Bayesian longitudinal growth model. This new structure allows users to correct for measurement error and capture individual-level variation in growth trajectories and parameters.
3. To demonstrate the new method we applied it to a sample of tropical tree survey data from long-term monitoring sites at Barro Colorado Island. We were able to reduce estimated error in size and growth, and extract individual-and population-level growth parameter estimates. We used simulation to evaluate the ability of the new method to improve estimates of growth rate and size, and estimate individual and species-level parameters. Our method substantially improved RMSE for growth by an average of 61% compared to existing approaches using pairwise differences; and reduced RMSE in estimated size RMSE compared to ‘observed’ values in simulated data. Better numerical integration methods (Runge-Kutta 4th order in comparison to Euler and midpoint) provided better estimates of parameters, but did not improve the estimation of size and growth. The choice of a positive growth function eliminated all negative increments without data exclusion.
4. Overall, this study shows how we can gain new and improved insights on growth, using repeat forest surveys. Our new method offers improved biomass dynamics estimation through reduced error in sizes over time, coupled with novel information about within-species variation in growth behaviour that is inaccessible with species average models, such as individual parameters for the growth function which allows for relationships between parameters to be considered for the first time.

## 1 Introduction

Repeat size survey data are used to estimate growth for many thousands of tree species across hundreds of long-term plots all over the world, in order to manage resources, estimate biomass, and model population dynamics (Condit, 1998; Anderson-Teixeira et al., 2015). Accurate growth estimates are essential as the growth behaviour across a wide variety of species is used to inform management policy and resource extraction, including estimates of carbon sequestration (Paniagua-Ramirez et al., 2021), ecosystem dynamics (Falster et al., 2018), regrowth (Lussetti et al., 2019), and logging yields (O’Hehir et al., 2000). Despite common use, the models built on repeat surveys from forest plots struggle to account for measurement error and tend to under-use the longitudinal structure. Repeat surveys are also used in other parts of ecology such as for modelling growth among fish (Hamel et al., 2014), where similar problems exist. In this paper we focus on situations where growth is primarily size-dependent, i.e. where the rate of change in size

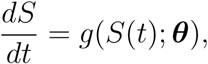

is determined by an individuals current size and some set of parameters ***θ***. The growth function *g* is chosen based on the data and desired features, and we consider ***θ*** to be random, taking different values for different individuals. Size-dependent growth functions have a long history in forest dynamics (Canham et al., 2004; Herault et al., 2011; Bhandari et al., 2021), as size is more important to growth than age and often age is not available anyway. This is the case for tropical tree species which do not have consistent annual rings (Condit, 1998), forests with episodic rainfall (Haines et al., 2018), or forests where the resources are not available to collect tree ring records due to the labour intensity (Anderson-Teixeira et al., 2015). While application of size-dependent analyses is common, existing methods for estimating growth mostly do not account for two key sources of variation, leading to biased estimates.

The first source of variation we will account for is measurement error in size estimates. While the true size of individual *i* at time *j* might be *S_i,j_*, we actually observe *s_i.j_* where:

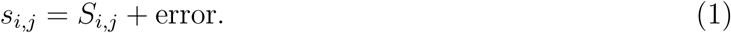

If we were to estimate growth of an individual in the time interval *t_j_* to *t_j_*_+1_ using a pairwise difference in observed sizes, *s_i,j_*_+1_ − *s_i,j_* (the most common current approach), we often introduce a lot of error into growth estimates, sometimes leading to nonsensical values. For example, in the simulated data of Figure 1, an unfortunate combination of measurement errors at years 7 and 8 (Figure 1a) led to a negative growth estimate at about 10cm (Figure 1b), for what is actually a fast growing plant, approximately doubling in size every two years. Negative growth estimates are easy to detect, but positive growth increments can also be errors and are much more difficult to spot. A longitudinal model offers the chance to correct those as well based on the surrounding information.

**Figure 1:**
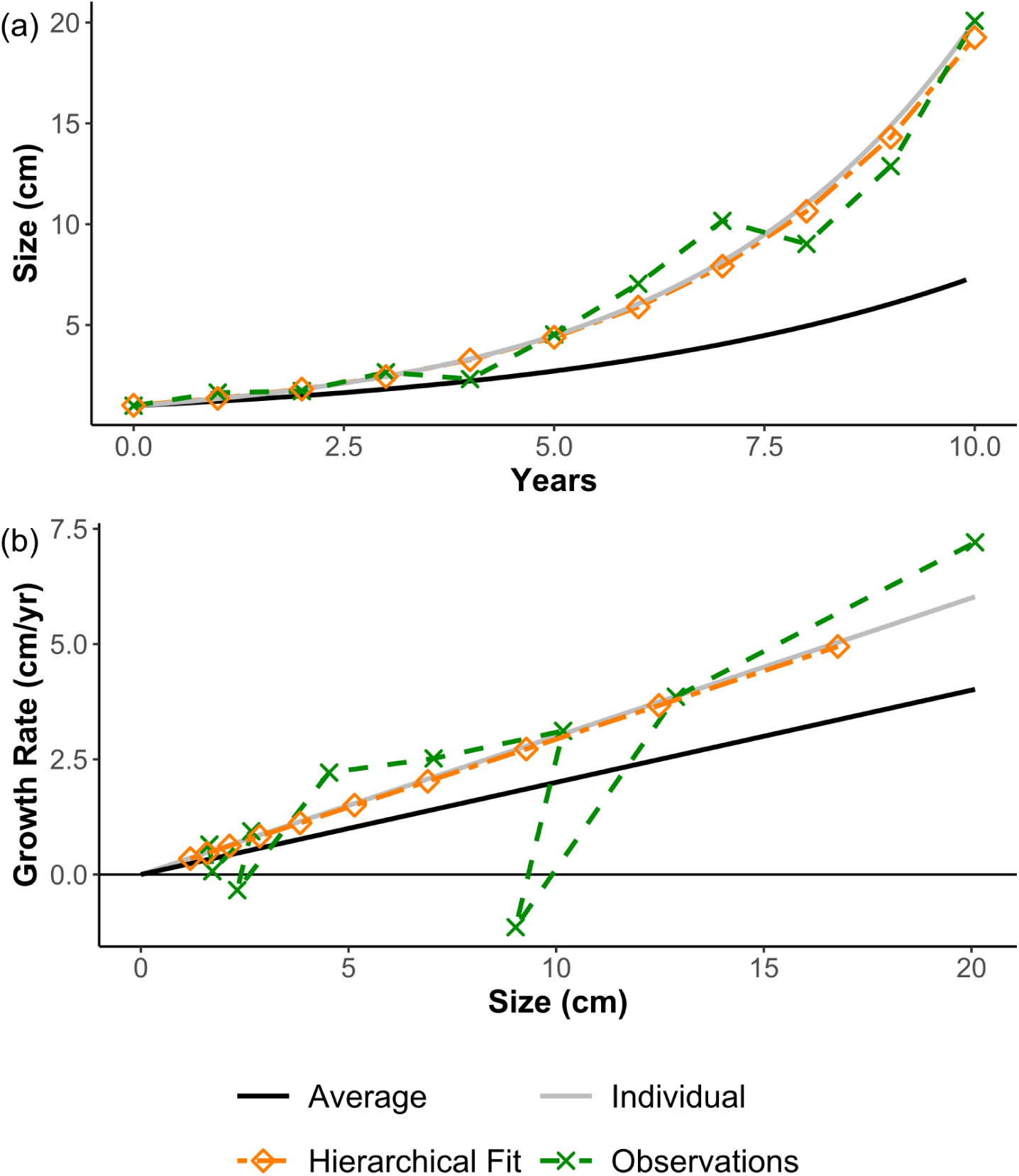
Example of measurement error and the impact on accuracy of growth measurements from simulated data with growth function *g*(*S*(*t*); *β*) = *βS*(*t*). **(a)** Simulated sizes of an individual with higher growth rate (*β* = 0.3) than the population average (*β* = 0.2). There are ‘observed’ sizes that include measurement error and corrected sizes after a hierarchical model has been fit. **(b)** A comparison between the true growth curves and the observed growth rates showing incidents of under-and over-estimated growth including negative increments, and the corrected estimates of growth from the fitted model.

A second source of variation we will account for is due to differences in growth trajectory across individuals within a species. In Figure 1, our example individual is growing faster than average, and attempting to describe the individual trajectory with the population average would introduce more error. While it is widely acknowledged that individuals within a tree species can vary greatly in their growth rates and trajectories, most modelling efforts still fit average models to a species, which neglects this variation.

In previous work, Aubry-Kientz et al. (2015), Bhandari et al. (2021), and Ganivet et al. (2017) modelled species-level growth trajectories with a single error term, but doing so conflates individual variation and measurement error. In such situations we cannot disentangle the two, and we would very much like to as individual variation within a species is relevant to understanding forest

dynamics (Falster et al., 2018), mortality (Camac et al., 2018), and trait distribution behaviour (Aubry-Kientz et al., 2015; Rüger et al., 2012). Size-dependent models have been used before by Ganivet et al. (2017) and Herault et al. (2011) but neither encode individual variation or longitudinal structure. A hierarchical model with individual levels was built in Clark et al. (2007), but is a Bayesian hierarchical equivalent of a linear mixed effects model for size with individual and year effects. The growth function in Clark et al. (2007) is constant conditioned on the year, rather than a size-dependent non-linear differential equation that we seek to use. Britton et al. (2007) and Hamel et al. (2014) used pairs of measurements, rather than a series of repeated measures, to parameterise a von Bertalanffy function for growth which does not use the longitudinal structure, while Canham et al. (2006) defines a function for the growth of trees which has parameters defined at the species level but without accounting for individual variation. Herault et al. (2011) has a species-level model equivalent to what we want to do, but fits growth functions to pairwise difference data and does not encode longitudinal structure or really access individual variation outside of proposing a quantile envelope around the average. Donnet et al. (2010) used a stochastic differential equation to model growth in chickens, with a single error term, which cannot distinguish between individual-level variation and measurement error. To the best of our knowledge there has not been previously been work using hierarchical models to disentangle measurement error from individual-level variation in size-dependent growth models.

Of the prior efforts to account for measurement error, the simplest is to exclude growth intervals if the estimate produced is too extreme which requires ad hoc exclusion criteria (Graham et al., 2021; Ganivet et al., 2017; Kenfack et al., 2014). We would prefer to correct for measurement error by including it in the model and exploiting auto-correlation instead of relying on data exclusion in pre-processing that involve chosen thresholds. Rüger et al. (2011) also builds a size-dependent model for errors based on the same short-interval repeat measurement data, and incorporates that into the hierarchical Bayesian model to estimate the error parameters, which we will detail below.

In this paper we develop a hierarchical, size-dependent growth model which captures individual-level variation in growth separate to measurement error, thereby overcoming some of the limitations in previous work. A hierarchical model lets us encode error and individual differences as separate sources of variation, thereby partitioning their impact on growth. Figure 1a demonstrates how such a structure might reduce error estimating size – and hence growth – while also capturing individual-level variation within a population. To demonstrate the potential of this new method on real data, we fit the model to long-term forest data from Barro Colorado Island. We then use simulation (i.e. data with a known truth) to evaluate the method’s effectiveness in estimating parameters.

## 2 Proposed approach

We use the mathematical notation given in Table 1 throughout this paper. The primary convention to note is the use of capital *S* for true sizes, *S̑* for estimates given by the chosen method, and lower case *s* for observations.

**Table 1:** Mathematical notation.

| Symbol | Interpretation |
| --- | --- |
| $S_{i,j}$ | True size from individual $i$ at time $j$. |
| $\hat{S}_{i,j}$ | Estimated size from individual $i$ at time $j$. |
| $s_{i,j}$ | Observed size from individual $i$ at time $j$. |
| $t_{i,j}$ | Time of census $j$ for individual $i$. |
| $g$ | Growth rate function chosen based on desired features. |
| $\Delta S_{i,j}$ | True growth increment for individual $i$ between times $t_{i,j}$ and $t_{i,j+1}$. |
| $\widehat{\Delta S}_{i,j}$ | Estimated change in size for individual $i$ over the interval between observations $j$ and $j + 1$, numerical integral approximating $\Delta S$. |
| $\Delta s_{i,j}$ | Pairwise difference in observed sizes given by $s_{i,j+1} - s_{i,j}$. |
| $\sigma_e$ | Global error parameter. |
| $\beta_i$ | An individual-level parameter which we treat as random. |
| $\mu_\beta$ | A population-level mean hyper-parameter for the individual-level parameter $\beta$. |

We have built a hierarchical Bayesian longitudinal model that leverages the autocorrelation structure of repeat measurement data in order to separate individual variation from measurement error. We assume individual trees follow a smooth growth trajectory that is reasonably approximated by our chosen growth function. By connecting a sequence of points, we can exploit the auto-correlation structure of repeat survey data to reduce measurement error. In doing so we assume each individual follows their own trajectory within the population distribution which means we can explore the within-population distribution structure. The individual growth function parameters are random parameters from a population-level distribution, that has its own random hyper-parameters, informed by the choice of prior distributions.

### 2.1 Growth model

We have repeat observations of size over time, *s_i,j_* for individual *i* = 1*, . . ., n* at time *t_j_* for *j* = 1*, . . ., m_i_*, which may vary across individuals. As we are looking at diameter measurement forest data we consider size to be diameter at breast height (DBH) in centimetres (Condit, 1998).

We assume these are measured with error and distributed as

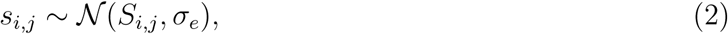

where *S_i,j_* is the true size and *σ_e_* is the measurement error standard deviation, independently for each *i, j*. The assumption of normality in this modelling framework needs to be flexible as Rüger et al. (2011) shows that we have non-normal error in the data, which we investigate in simulation. We assume changes in size across sampling times are a function of an *a priori* defined growth function *g*(*S*(*t*); ***θ****_i_*) such that

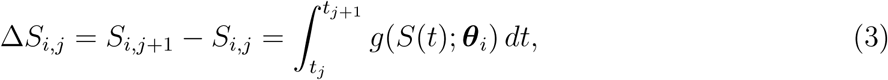

where ***θ****_i_* represents a vector of individual-level growth parameters.

Recall that we emphasised that our model accounts for two key sources of variation – measurement error and individual-level variation. Equation (2) is how we represent measurement error, and the parameter vector ***θ****_i_* in Equation (3) is how we represent individual-level variation in growth trajectory.

The growth function used in this paper is the Canham growth model (Canham et al., 2006), given By

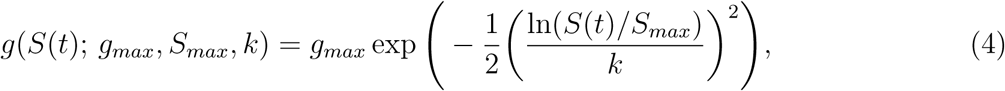

where *g_max_* is the maximum growth rate, *S_max_* is the size at which that growth rate occurs, and *k* controls the decay after the peak. We chose Equation (4) due to literature precedent and useful features. It is bounded above, has a hump shape that predicts a lower absolute diameter growth rate for small sizes. With different parameter combinations Equation (4) can model increasing, decreasing, peaked, or stable growth rates. The model has biologically interpretable parameters and is bounded above by *g_max_*, so does not predict infinite growth like an exponential function. In addition, Herault et al. (2011) found it to perform the best of the growth models that were implemented to capture the growth behaviour of tropical trees. Further details about the behaviour of the Canham model are in the supplmentary material.

### 2.2 Estimation

We elected to use MCMC because it is a commonly used method that is known to flexibly estimate posterior distributions.

At the individual and hyper-parameters level, we use 95% credible intervals (CIs) from the posterior distribution samples to summarise paramter distributions. While there is discussion (Kruschke, 2014; McElreath, 2020) about the most effective interval construction method – and the best choice of quantile limits – we chose a common and accessible (to people with minimal Bayesian statistics) approach of using the central 95% credible interval rather than a highest posterior density interval. We also looked at density plots of the errors for hyper-parameters to see if estimation is well behaved or shows evidence of pathology.

For all models we assume

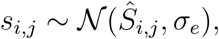

independently for each *i, j*, where *σ_e_* is a global error term. At the individual level, we fit the first size as a parameter with prior centred at the observed first size

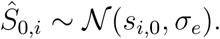

The priors for each parameter are detailed in Table 2. The log-normal distributions are chosen in line with Herault et al. (2011), but also because Equation (4) is parameterised in such a way that non-negative values give the correct functional form. The mean and standard deviation hyper-parameters for the log-normal distribution also have useful interpretation in the context of the population.

**Table 2:** Priors for parameters across the full hierarchical model with individual parameters, and the species-level model based on (Herault et al. 2011)

| Model | Parameter | Prior |
| --- | --- | --- |
| Canham with individual parameters $g_{max}$, $S_{max}$, $k$ | $g_{max,i}$ | $\ln \mathcal{N}(\mu_{\ln(g_{max})}, \sigma_{\ln(g_{max})})$ |
| | $S_{max,i}$ | $\ln \mathcal{N}(\mu_{\ln(S_{max})}, \sigma_{\ln(S_{max})})$ |
| | $k_i$ | $\ln \mathcal{N}(\mu_{\ln(k)}, \sigma_{\ln(k)})$ |
| | $\mu_{\ln(g_{max})}$ | $\mathcal{N}(0, 1)$ |
| | $0 < \sigma_{\ln(g_{max})}$ | $Cauchy(0, 1)$ |
| | $\mu_{\ln(S_{max})}$ | $\mathcal{N}(0, 1)$ |
| | $0 < \sigma_{\ln(S_{max})}$ | $Cauchy(0, 1)$ |
| | $\mu_{\ln(k)}$ | $\mathcal{N}(0, 1)$ |
| | $0 < \sigma_{\ln(k)}$ | $Cauchy(0, 1)$ |
| | $0 < \sigma_e$ | $Cauchy(0, 2)$ |
| Canham with species parameters $g_{max}$, $S_{max}$, $k$ | $g_{max}$ | $\ln \mathcal{N}(0, 5)$ |
| | $S_{max}$ | $\ln \mathcal{N}(0, 5)$ |
| | $k$ | $\ln \mathcal{N}(0, 5)$ |
| | $0 < \sigma_e$ | $Cauchy(0, 1)$ |

#### 2.2.1 Numerical integration

In order to fit the statistical model to sizes over time we need to integrate the growth function between two time points to calculate

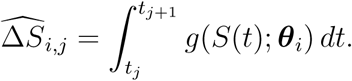

While for some growth functions an analytic solution exists, there is none for Equation (4), or in general for others. Hence, to approximate the integral we need to use some form of numerical integration.

We implemented a 4^th^-order Runge-Kutta algorithm, as it is known to be more accurate than simpler alternatives such as Euler or midpoint methods(Butcher, 2016), but tractable within the available computation time. As error does arise from numerical methods, a comparison of three can be found in the supplementary material. The comparison shows that simpler integration methods tend to give biased estimates of model parameters to compensate for the higher errors that result from worse integration. The 4^th^-order Runge-Kutta algorithm overcomes the bias and returns accurate parameter estimates. Accuracy of the integration was further improved by taking multiple substeps of 1 year, to step the growth function across the desired time interval (5 years in the data). Going from an observation interval of 5 years to five sub-steps of 1 year increases the required calculation at each point.

### 2.3 Model evaluation

As a quantitative sanity check on the predicted sizes in real-world data we use the *R*^2^ statistic to compare observed and estimated sizes over time. We calculate *R*^2^ for each individual in the real world example, and for the overall sample.

In order to evaluate how well our method reduces the impact of measurement error on growth, we used simulated data with known parameters and sizes to understand how well error correction behaved. For the simulated data we first added measurement error to ‘true’ sizes, then fit the model and compared the estimated sizes and modeled growth for each interval to the true values using the root mean squared error (RMSE) statistic. We first used normally distributed error, then a mixed distribution error model based on Rüger et al. (2011). To show evidence that our system reduces error we compare the RMSE of the estimated increment values Δ*S̑* = *s̑_i,j_*_+1_ − *s̑_i,j_* to the RMSE of growth estimates given by the pairwise difference

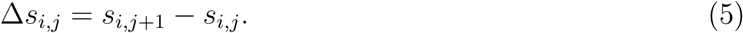

We also use the RMSE at the level of an individual size to compare the modelled size *S̑_i,j_* to the observed size with measurement error *s_i,j_* .

We have a couple of ways to evaluate the growth parameter estimation. We use scatter plots and RMSE of *θ̑_i_* − *θ_i_,* where *θ_i_* is the true value of that growth parameter for the individual and *θ̑_i_* is the estimate. The closer the difference is to 0, the better an estimate. The difference also allows us to check whether our estimation is biased: if the centre of the differences for the whole sample is 0 then we can be reasonably confident that our method is unbiased.

In line with Gelman et al. (2021), we use the *Ȓ* statistics to evaluate MCMC chain convergence for a given parameter, which compares the between-chain variance and within-chain variance. Histograms of *Ȓ* are given in the supplementary material where relevant. We consider the chains to be poorly behaved if there are many *Ȓ >* 1.05, a more liberal limit chosen to encompass that the data we use can be messy for particular individuals, and we have many individual-level parameters to fit. Traceplots are also provided and allow for additional visual inspection of convergence behaviour.

To test the ability for our models to predict out of sample sizes we did one-step-ahead estimation. We built two additional hierarchical models to the first five observations: one with the Canham growth function and one with a constant growth function which estimates an individual average growth rate. The Canham model performed better than using individual average growth. Details are in the supplementary material.

### 2.4 Implementation

The model was fitted using MCMC using the RStan package (Stan Development Team, 2019), which serves as an interface to the Bayesian modelling system Stan (Stan Development Team, 2022). We used R version and 4.3.1 (R Core Team, 2018). Additional computational resources were provided by Katana (2010) at the University of New South Wales. Code for this paper is available at https://doi.org/10.5281/zenodo.14048054 (O’Brien et al., 2024), which includes model fitting, example data, and analysis code, but not the automation code used to run models on Katana.

Runtime for these non-linear models can vary considerably with the low end being 24-48 hours for 400 individuals, and the high end exceeding 200 hours for the slowest chain. Memory requirements are on the order of 4-8GB of RAM for the 400 individuals in the real data. Computation was much faster and less intensive for individual simulations with 50 individuals, details are provided in the supplementary material. With these computational requirements it is possible to run models on a laptop, subject to time constraints. Longer running models make more consistent computation infrastructure such as a server desirable. Large sample sizes or many replicates in the case of simulation indicate that a typical PC may not be sufficient.

## 3 Case Study: Dynamics of *Garcinia recondita* in 50 ha plot at Barro Colorado Island, Panama

To demonstrate the method on real data we took a sample from the species *Garcinia recondita* in the Barro Colorado Island (BCI) dataset (Condit et al., 2019). The sample also allowed us to extract reasonable parameter estimates, which we use to inform the simulations in following section.

### 3.1 Data

The forest plot at BCI has had a census of diameter at breast height (DBH) for all plant stems over 1 cm in diameter every 5 years since 1985 (Condit et al., 2019). *G. recondita* was chosen for having a large enough population size to take a sample of 400, and for being among a list of tree species of particular interest to us: a shade tolerant species that can survive well in low light environments.

We filtered the dataset to select a sampling frame by including only records with the following criteria:

- observations at 1.3m above the ground,
- taken after the change in measurement precision from 5mm to 1mm in 1990,
- verified matching of stem and tree IDs, and
- six observations available for the individual (the maximum possible in the observation period).

Excluding individuals without the maximum number of observations excludes both those recruited after the first census, and those that died during the observation period. We did not exclude data with apparently negative growth increments between intervals as the choice of a positive growth function will enforce positive increments. There are 2,778 *G. recondita* trees in the BCI data satisfying these criteria, and we built a sampling frame from the tree IDs. For this study, we took a simple random sample without replacement of 400 individuals, giving us 400 sets of 6 observations, with approximately 5 years between them.

### 3.2 Results

The trace plots and a histogram of *Ȓ* values are given in the supplementary material. Model convergence was good, both individual-level and species-level chains behaved well. The model took 26 hours to run and used 4GB of RAM.

#### 3.2.1 Estimated and observed sizes

Figure 2 gives five examples of fitted and observed sizes for individuals, demonstrating different types of behaviour. The *R*^2^ statistic, calculated as the squared correlation of observed and estimated sizes, was used as a quantitative tool to analyse goodness of fit. Panel (a) shows a close match between observed and fitted sizes, with steady growth increments across the duration, and an extremely high *R*^2^ value of 0.996. Panel (b) shows more size correction, with less overall growth – and a more consistent growth rate – in the observation interval. Panel (c) shows a close match between the observed and estimated values with a tree growing past its *S_max_*, demonstrating the potential to fit a hump-shaped relationship between growth rate and size. In a plot of size versus year, such trajectories have an inflection point and asymptote. The peak growth rate for this individual is close to 1cm*/yr* between 2005 and 2010, which is very high when compared to the rest of the sample. Panels (a) and (b) have quite similar fitted behaviour, though the former has higher growth than the latter and reaches a much greater size. In Panel (d) we can see an indi-vidual with very low growth across the observation period, and very noisy data including negative growth increments. The overall change in size for this individual is very small, so the negative increments look more than they are compared to the other individuals. The overall sample *R*^2^ is 0.995, showing extremely strong agreement between estimated and observed sizes, and individuals with low *R*^2^ tended to have low overall growth and negative increments. More details are available in the supplementary material.

**Figure 2:**
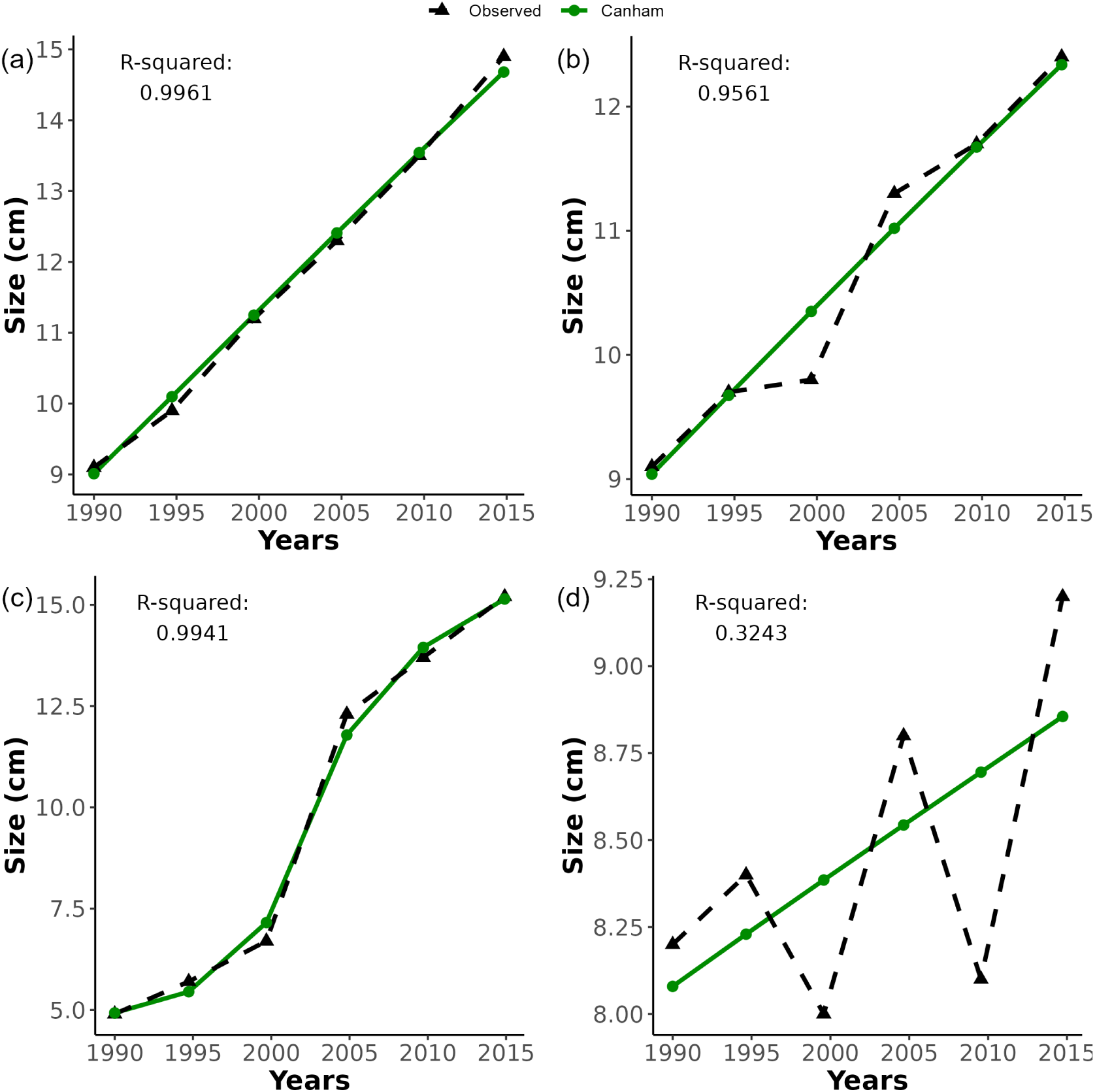
Examples of individual trees from *G. recondita* showing the observed sizes and the smoothed size tra-jectories from the fitted Canham model. **(a)** and **(c)** show close fits to observed sizes going past a growth peak. **(b)** shows smoothing of some sizes. **(d)** shows smoothing of both the negative increments, and subsequent large positive increments.

Our model completely eliminated the negative growth increments in Panel (d) by applying a positive growth function in Equation (4), but also reigned in the extreme positive increments that come after them. If we consider the observed increment in Panel (d) between 2005 and 2010, we have 0.8cm over 5 years, or 0.16cm*/*yr, which is not an extreme growth rate in the overall sample when we compare it to the individuals in Panels (b) and (c) for example. As such, it would be difficult to detect that the 2005-2010 growth increment of 0.8 cm in Panel (d) is likely an error without fitting the individual trajectory. By contrast, the estimated growth increment is 0.16 cm.

#### 3.2.2 Fitted growth functions

Our model fitted function parameters to each individual, producing different growth curves. Figure 3 gives a visual comparison between individuals and species-level average trajectories from both a species-level model. The species-level model was fit in line with Herault et al. (2011) as a function rather than a differential equation, with data constituted of independent, annualised pair-wise size differences without a longitudinal structure:

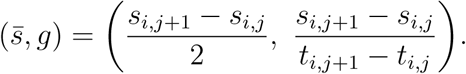

**Figure 3:**
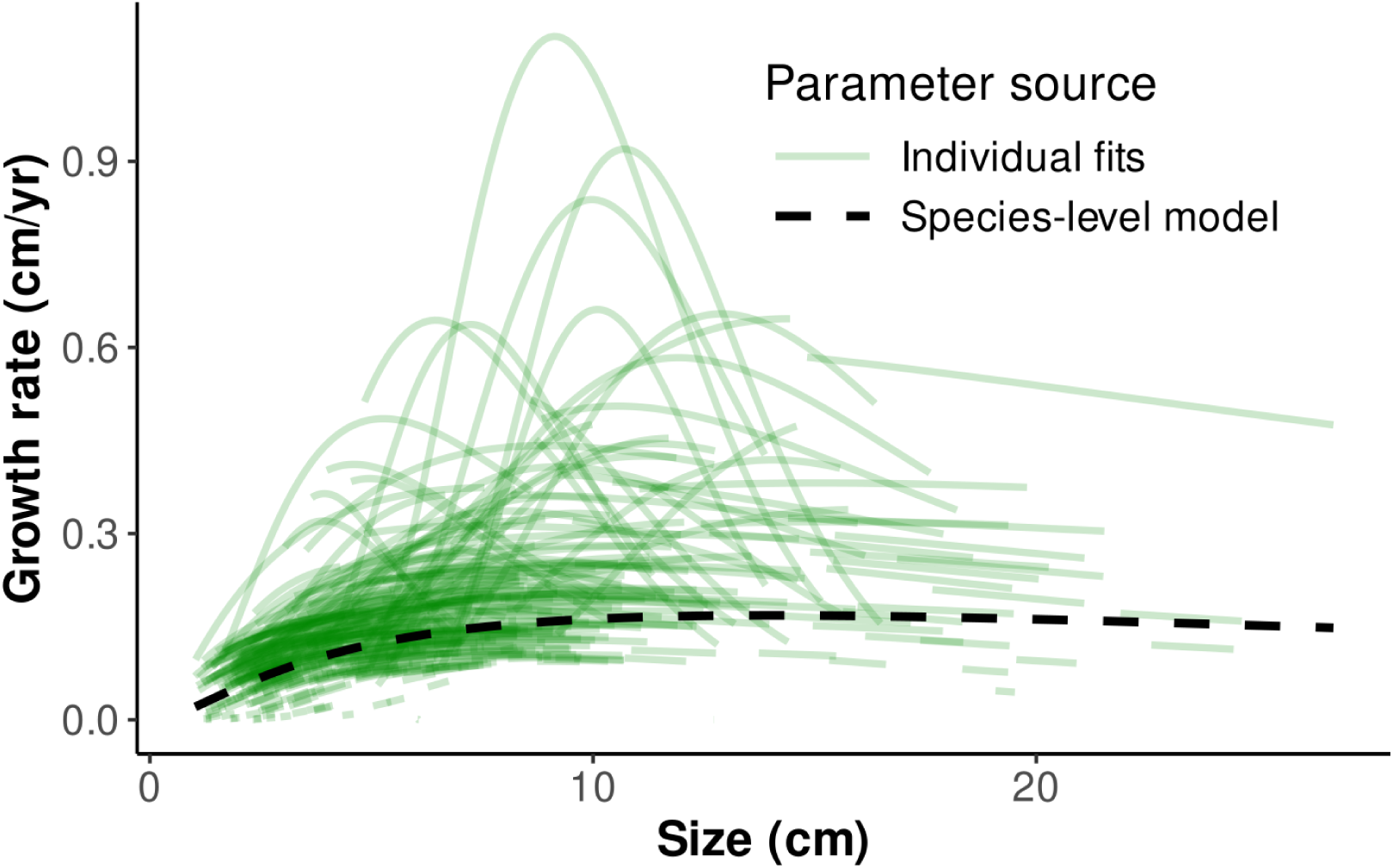
Individual growth trajectories from *G. recondita* (green lines) show considerable variation in growth trajectory that is not captured by a species-level model as in Herault et al. 2011). Note in particular that individuals tend to display a steep growth decline at large sizes that is not apparent from the species-level model.

Individual trajectories depart from this species-level average, for example the species-level model estimates the max growth rate to be 0.168 cm/yr, while 37 of the 400 individuals (9.25%) had *g_max_* in excess of twice this value. As well as the maximum, the shape of individual curves differs from the species level fit. Most large individuals show a decline in DBH growth at large sizes, a trend that is not evident in the species-level model but is clearly possible with the Canham growth function.

Plotting the estimated growth trajectories in Figure 3 allows us to get an idea of any trends across the population. In this case we do see a general inclination to sustained modest growth rates across sizes. About a dozen function pieces show very peaked high max growth behaviour, while the bulk of the 400 individuals fall into a more consistent band of slower growth with a slight decline for larger sizes.

In Figure 3 there is a cluster of individuals with near-zero growth across the entire observation period, which can be seen close to the *x*-axis. These are real individuals, and provide a statistical challenge as parameter estimation struggles when no or little change is observed. We can reasonably anticipate that parameter estimates for individuals where so little of a growth function is seen are unreliable even if the growth and size estimates are believable when compared to observed sizes over time.

When we compare the species-level model in Figure 3 to the individual pieces and median trajectories, we see that the species-level model has a later peak with *S_max_* = 14.0 compared to the median *S_max_* = 8.81. Furthermore, the species-level model does not decline in growth for the larger observed sizes as fast as the individual pieces in that part of the distribution. A lot of the sample sizes are smaller than 10 cm, with very few trees larger than 15 cm. A function fit to piecewise differences will be more strongly influenced by the density of data at the small end of the distribution than the few individuals with larger sizes as both *k* and *S_max_* are influenced by how fast accelleration is at small sizes. Due to the high mortality of trees, we expect few of those small individuals to grow to maturity, so a population level trajectory that includes them may not work well for the survivors.

Figure 3 demonstrates that an average trajectory alone is insufficient to describe behaviour within a species, particularly when we take into account the size distribution. Only by fitting individual trajectories are we able to ease apart the diversity in growth behaviour across the sample. We give greater detail on the individual parameter distributions in the supplementary material.

#### 3.2.3 Population-level parameters

At the species level we get point and 95% CI estimates given in Table 3. The species-level averages for max growth and the size at which it occurs are 0.16cm/yr and 8.0cm respectively. The average for the spread parameter *k* is more difficult to interpret, but indicates that at three times the peak size (24cm) DBH is increasing at 37% of the peak rate.

**Table 3:** Population level hyper-parameter estimates from the *G. recondita* sample to 2 sig. figures.

| Parameter | Estimate | 95% CI |
| --- | --- | --- |
| $\mu_{\ln(g_{max})}$ ($\exp(\mu_{\ln(g_{max})})$) | -1.85 (0.16 cm/yr) | (-1.96, -1.74) |
| $\sigma_{\ln(g_{max})}$ | 0.64 | (0.56, 0.71) |
| $\mu_{\ln(S_{max})}$ ($\exp(\mu_{\ln(S_{max})})$) | 2.08 (8.0 cm) | (1.96, 2.20) |
| $\sigma_{\ln(S_{max})}$ | 0.47 | (0.36, 0.58) |
| $\mu_{\ln(k)}$ ($\exp(\mu_{\ln(k)})$) | -0.24 (0.78) | (-0.40, -0.061) |
| $\sigma_{\ln(k)}$ | 0.64 | (0.51, 0.78) |

## 4 Simulation

The case study above shows the potential of the new method to reveal novel information about growth in tree communities. The question remains: are these estimates any good? It’s impossible to know using field data, as the truth is unknown. Instead, we use simulation to assess how accurately we can estimate size, growth, and function parameters using our new methods. In simulation we know the underlying true values and behaviour and are thus able to compare model estimates directly to the truth which is inaccessible in real observations. We want to look at reduction in error for size and growth estimates compared to observations, whether we get reasonable estimates of individual parameters, and whether we can reliably estimate population-level hyperparameters.

To achieve this, we simulated 50 individuals growing according to Equation (4), with individual parameters sampled from a trivariate log-normal distribution with the means and covariances set to their sample estimates based on our fit to *G. recondita* from Section 3. We simulated initial sizes (*S*_0_) from log N (2, 1), with values restricted to between 1 cm and 15 cm to allow for growth as we do not see *G. recondita* smaller than 1 cm or larger than 25 cm. The maximum *S*_0_ maintained the upper bound of 25 cm for plausible amounts of growth over the observation period. More detail is given in the supplementary material including exclusion criteria as not all parameter combinations produce plausible behaviour. From each individual’s life trajectory we took 6 size measurements, 5 years apart, to match the survey structure of Barro Colorado Island. These 300 true sizes formed the basis of simulations with measurement error.

For these 50 individuals with a known true growth trajectory, we then simulated ≥ 1000 realisations of the measurement process, adding noise to the data, as would occur with any field sample.

Simulated sizes with error become ‘observations’ and are the basis of a particular model fit. This simulated data allows us to assess three aspects of model performance: accuracy estimating true size and growth, the ability to recover individual parameters, and the ability recover species hyper parameters.

### Size and growth

For growth increments, we compare our model estimates (Δ*S_i,j_* = *S̑_i,j_*_+1_ − *S̑_i,j_*) and observed pair-wise differences (Δ*s_i,j_* = *s_i,j_*_+1_ − *s_i,j_*), in terms of how well they estimate true growth (Δ*S_i,j_* = *S_i,j_*_+1_ − *S_i,j_*). For size, we compare our model estimates (*S_i,j_*) to observed sizes (*s_i,j_*), as estimates of the true size *S_i,j_* . We use root mean squared error (RMSE) as our measure of predictive acccuracy, and consider the mean percentage reduction in error between observed data and the values fitted by the model.

### Individual growth parameters

For our hierarchical model, we look at the mean proportion of posterior central 95% credible intervals that cover the true individual-level values. The proportion is calculated for each of the simulations and each of the three individual-level parameters. We also look at RMSE and the distribution of posterior estimates for individuals to check for evidence of bias and size-dependent error.

### Species growth hyperparameters

We look at posterior 95% CI coverage of the true parameter values for the hierarchical model, as well as examining the distribution of estimate errors *θ̑* − *θ*.

In the simulations, we considered two alternative error processes:

1. **Normal error**: A size-independent normally distributed error process

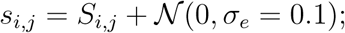

2. **Rüger error**: A process with both size-dependence and catastrophic error (Rüger et al., 2011), given by

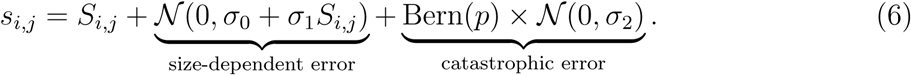

The second error model has both a realistic error simulation that is not properly matched with the assumed normal distribution in the Bayesian model, and higher overall error than the first model due to the size-independent catastrophic errors. Parameter values were set at *σ*_0_ = 0.0927cm, *σ*_1_ = 0.00038, *p* = 0.0276, and *σ*_2_ = 2.56cm, best fit estimates from Rüger et al. (2011). For a 12cm individual, the size-dependent error has a standard deviation of 0.09726cm. Combined with the second larger normal distribution (catastrophic error) that occurs with 2.76% probability in Equation (6), negative growth increments and large errors are more common for the Rüger error compared to the N (0, 0.1), while the 97.24% non-catastrophic errors are of comparable size.

To guarantee a minimum sample size of 1000, allowing for pathologies in the chain convergence and computation time overrun, we ran 1100 independent simulations from each error process. We extracted max *Ȓ* values and number of divergent transitions from model output and excluded any with more than 50 divergent transitions and a max *Ȓ >* 1.05. These are very permissive thresholds, chosen as we know the true values of sizes and parameters, and want to see what model performance looks like even with moderate issues.

### 4.1 Results: New method improves estimate of size and growth

In the first set of simulations we used normally distributed additive error. Of the 1100 simulations, 1095 finished within the max time and five were excluded on *Ȓ* and divergent transition grounds. We used all of the remaining 1090 for further analysis, with a mean runtime of 52 minutes.

*Size and growth* Error scatter plots for an example simulation in Figure 4 show what reduction in error looks like at the level of measurement. On average we saw 29.4% reduction in size error and 61.1% reduction in growth error compared to ‘observed’ sizes and growth increments. Figure 4(c) and (d) give histograms of the percentage reduction across all simulations, which are symmetric and show that in no simulation did applying our method increase error.

**Figure 4:**
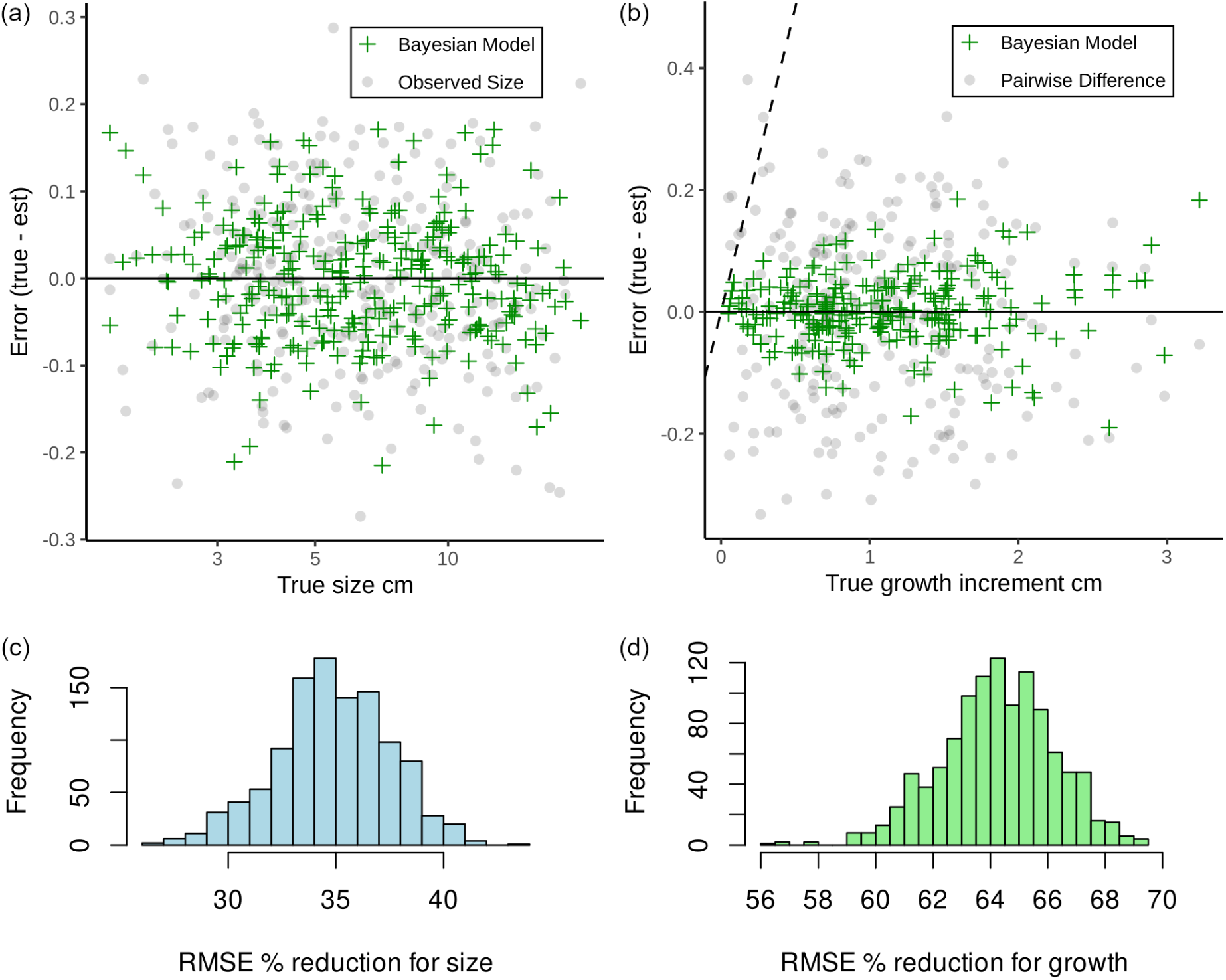
Error reduction in size and growth for simulations with 𝒩 (0, 0.1) added error. **(a)** shows errors against size in an example simulation for *si,j* observed sizes and *S̑i,j* fitted sizes. In this example we have a 28% reduction in size error. **(b)** gives the errors for growth increments and shows a 61% reduction in growth error for this example. Points above the diagonal dashed line are negative growth increments in the observed data, all of which are corrected without data loss in the fitted values. **(c)** and **(d)** show the distribution of size and growth RMSE error reduction across the N (0, 0.1) simulations, and that in no cases did the model increase error.

The difference between size and growth error reduction is a result of what we call parallel growth increments, where the change in size is the same, but the sizes are not, for example we might have

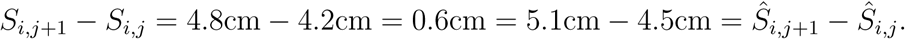

An example of reduction in error for size and growth can be seen in Figure 4(a) and (b) scatter plots of the true values *S_i,j_* against error from observed *s_i,j_* or estimated *S̑_i,j_* . The reduction is fairly symmetric across the sizes (aside from the negative values), and through the use of the longitudinal model at the individual level we are able to constrain errors which are not extreme in the whole population, but are extreme for that particular individual.

*Individual growth parameters* results are summarised in Table 5 and scatter plots in Figure 6 that compare estimated parameter errors to true values. For *g_max_* we have good coverage, avoid the shrinkage towards the mean which is typical of hierarchical models, and have no evidence of size-dependent bias. For *S_max_* there is no clear indication of size-dependent bias in Figure 6(b), however Panel (c) shows that higher values of *k* tend to be under-estimated by the model more than lower ones are over-estimated as would be expected, through shrinkage, in a hierarchical model. Overall, the estimated individual parameters do show a slight reduction in variance compared to the true values. The bias of a particular individual’s parameters is likely due to which part of the growth function is observed as individuals with similar parameters demonstrate different biases.

**Table 4:** RMSE across 1000 simulation for estimates of growth and size comparing model fit to observed size (with measurement error) and pairwise difference in sizes. The Bayesian method shows considerable improvement in estimates for growth, and more modest improvement in estimates for size.

| Error Type | Measure | Mean RMSE |  | Mean RMSE |
| --- | --- | --- | --- | --- |
|  |  | Obs. | Model Est. | % Reduction |
| $\mathcal{N}(0, 0.1)$ | Size | 0.10 | 0.0651 | 34.9% |
|  | Growth | 0.141 | 0.0504 | 64.2% |
| Rüger Eqn. (6) | Size | 0.121 | 0.065 | 46.2% |
|  | Growth | 0.171 | 0.050 | 70.4% |

**Table 5:** Individual-level parameter estimation for Canham simulations.

|  |  | Mean 95% CI | Mean CI | Mean |
| --- | --- | --- | --- | --- |
| Error Type | Par. | Coverage | Width | RMSE |
| $\mathcal{N}(0, 0.1)$ | $g_{max}$ | 94.2% | 0.122 | 0.323 |
| | $S_{max}$ | 94.0% | 7.83 | 11.0 |
| | $k$ | 92.0% | 1.10 | 1.78 |
| Rüger Eqn. | $g_{max}$ | 94.1% | 0.122 | 0.324 |
| | $S_{max}$ | 94.0% | 7.76 | 11.0 |
| | $k$ | 92.0% | 1.09 | 1.78 |

The individual with the least growth – No. 35 – has been highlighted in Figure 6 to demonstrate the impact that such a small change in size has on estimation. In Panel (a) the estimates consist of the values at *g_max,_*_35_ = 0.480 to the left of the red asterisk, but not those close to the 0 line which is a separate individual with *g_max_* = 0.481, indicating negative bias in estimating this parameter. For comparison *k*_35_ has a symmetric spread of error around the true value of 0.353.

For *S_max,_*_35_ = 9.88, the full spread above the red asterisk are the estimated values which indicates a lack of identifiability – multiple different parameter values which provide very similar observed behaviour (Talts et al., 2020), which is not clearly present for either *g_max_* or *k*.

*Species growth hyperparameters* results are summarised in Table 6, and visualisations of error distributions in Figure 5. For most parameters, we have coverage probabilities in excess of the 95% we would expect from a 95% CI in typical inference, while for *s*_ln(*S*_*_max_*_)_ the coverage probability is lower (92%), but still very good. In Figure 5 the estimates for *g_max_* mean and standard deviation, and the mean for *S_max_* show good behaviour with nice bell curves, though the curves are not centred at 0. The other parameters show less cooperative behaviour. For *S_max_*, while the mean parameter was well behaved *s*_ln(*S*_*_max_*_)_ has a positively skewed distribution of estimates centred above the true value, which aligns with the positive bias we see in the summary statistics. Estimation of *k* mean and standard deviation was spread out and inconsistent, with slight over-estimation of the average, and under-estimation of the spread.

**Figure 5:**
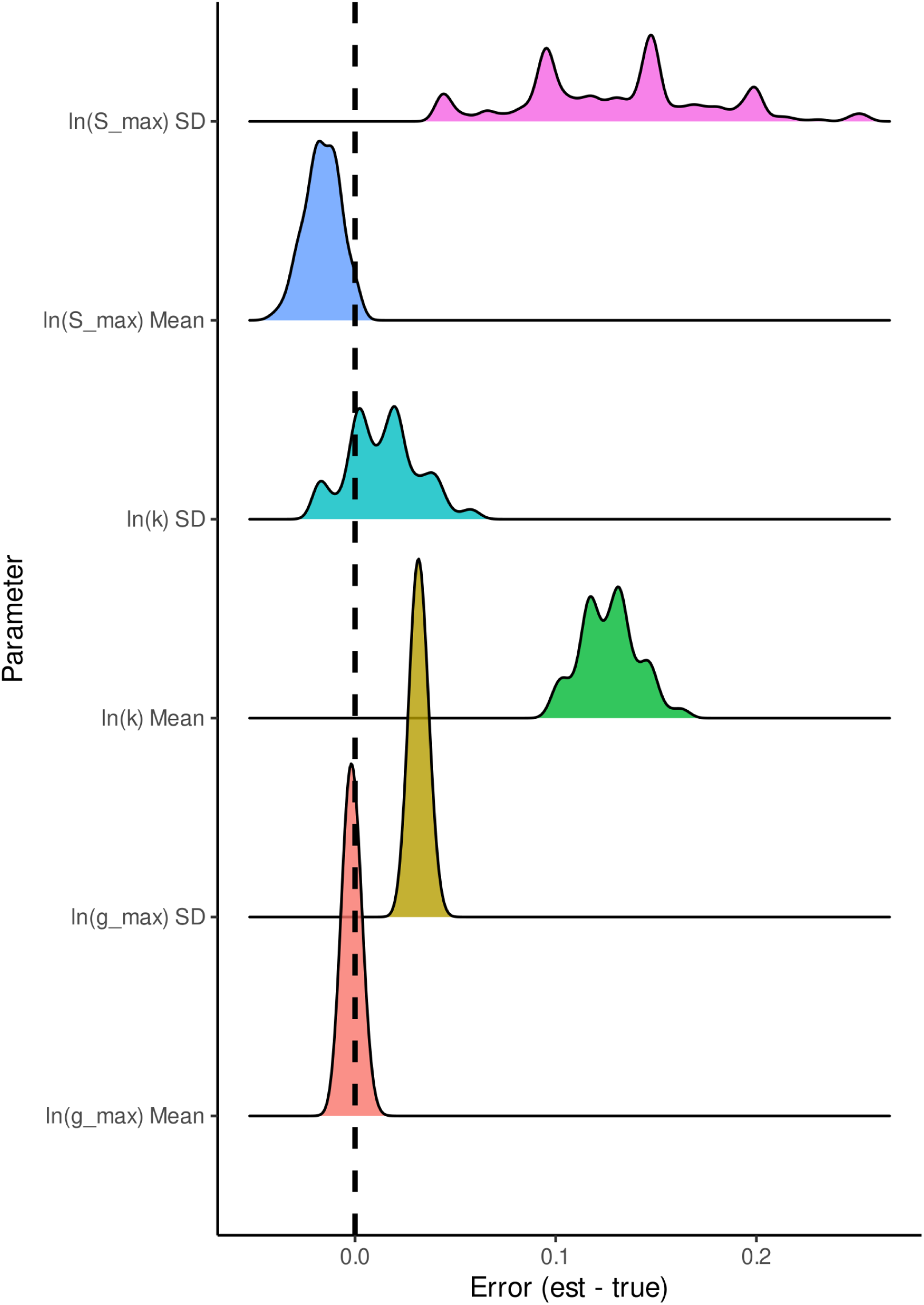
Density plot shows that estimation is better for some parameters (those for *gmax* for example) compared to others. The true values are in Table 6.

**Figure 6:**
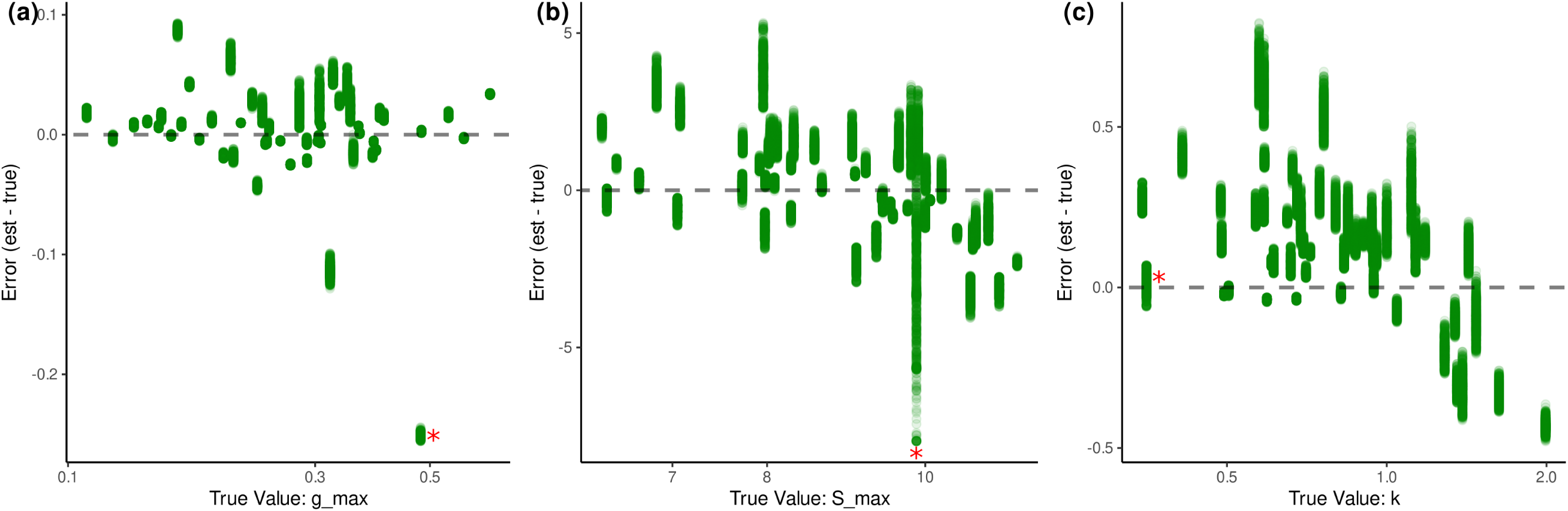
Scatter plots of individual parameter estimate errors with 𝒩(0, 0.1) as measurement error process. While *S_max_* and *k* show shrinkage towards the mean, *g_max_* does not. As the scatter is seemingly random for individuals with individuals with similar parameter we suspect that bias is related to what of the growth function is observed for a given individual. The red asterisk points to the estimates for individual with smallest total growth (0.32 cm) and demonstrate the difficulties that come with estimation under limited information.

**Table 6:**
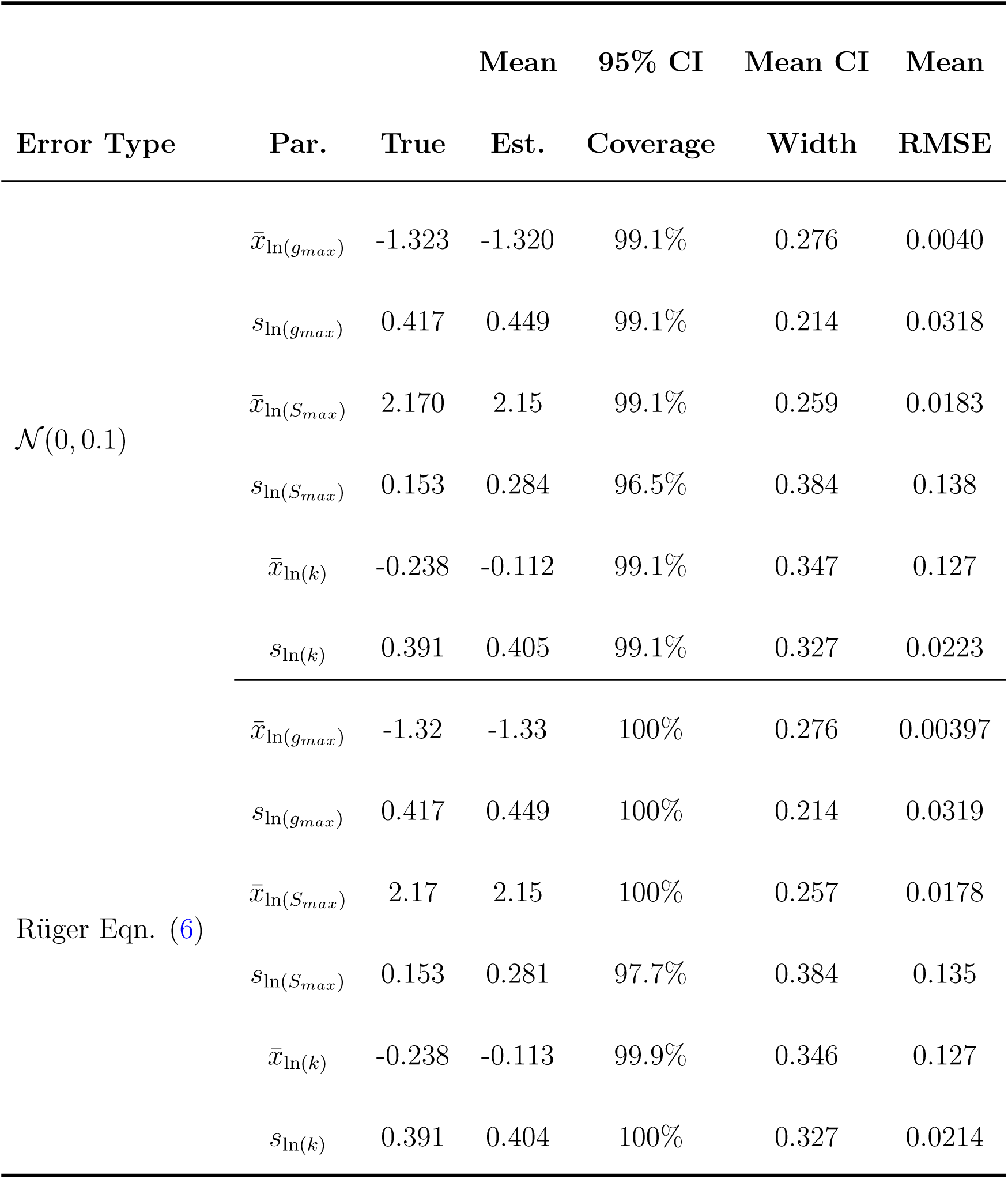
Population level hyper-parameter estimates for Canham simulations with different added error, with posterior central 95% credible intervals.

*Global error* estimation in the form of *σ_e_* had a mean estimate of 0.0925, compared to a true value of 0.1. The 95% CI mean width was 0.0192, and the coverage of 0.1 was 99.8% so while the error was slightly under-estimated it was not a huge problem.

*Rüger error model* results were very similar to those with the normally distributed error at the measurement, individual, and species levels as seen in Tables 4 and 6 despite the increase in overall error and occurrance of extreme errors. Estimation of *σ_e_* has a different interpretation as the true error distribution is not normal, but the posterior estimates had a mean of *σ_e_* = 0.0915 which matches the normal error estimates. Further comparisons can be found in the supplementary material. As the Equation (6) error generation process produces very similar results we can be comfortable using the N (0*, σ_e_*) likelihood for *G. recondita*.

## 5 Discussion

The use of repeat survey forest plot data to estimate growth behaviour has been plagued with measurement error problems (Condit, 1998) that we seek to solve by implementing a hierarchical Bayesian longitudinal model for sizes over time based on a chosen growth function. Errors in size can produce growth increment errors that under-or over-estimate size change, and for smaller sizes in particular such error can lead to biologically unlikely negative growth increments which are easy to detect, but have usually been dealt with by data exclusion. By dealing with measurement error we have opened up access to better estimates of size dynamics for the many long term forest plots reliant on repeat measurement surveys. Our new method also allows us to access new information in the form of individual parameter estimates and a richer picture of growth curve variation across a population. These improvements in size and growth estimates exist even in the context of an error process that is more complex than size-independent normal.

Although we have focused on trees here, the method is applicable to a very general class of problems. For example, Equation (4) is distinctly non-linear. The only requirements for applications outside tree growth dynamics are repeat measurement data with enough observations at the indi-vidual level to fit the relevant growth function, and a suitable differential equation to describe the dynamics. The method is of particular interest when the variation within a population is relevant information to analyse, or when there are repeat surveys involving non-trivial measurement error. As such, the method might also be deployed in other fields such as environmental processes or engineering where longitudinal data for smooth dynamical systems is available.

In this study, we were able to successfully fit the non-linear Canham model to example diameter at breast height (DBH) data from Barro Colorado Island. The data from *G. recondita* is very sparse (observations have 5 year time intervals) which makes estimation difficult. We see plausible sizes over time for individuals that are closely aligned with observation, but also eliminate negative increments without data loss in comparison to methods that require data exclusion (Graham et al., 2021; Ganivet et al., 2017; Kenfack et al., 2014) which would have reduced the already sparse dataset even further. By fitting the individual trajectories we achieve a lower predictive error compared to species level models such as Herault et al. (2011). Building in the longitudinal structure also improves the predictive error when compared to models that treat increments from the same individual as independent (Bhandari et al., 2021; Herault et al., 2011). Encoding individual variation of growth trajectories allows us to separate individual variation from measurement error, which in turn reduces the difference between estimated sizes and the observations as we are not forcing individuals to match the sample average. The next step with Barro Colorado Island data will be to apply this method to multiple species. We expect that the individual trajectories and parameterisations will give access to much richer information about individual variation within species, and as a result more ways to compare growth behaviour across species as well.

We were able to exploit the posterior estimates from *G. recondita* to build a parameter distribution for simulation which allowed us to constrain the behaviour of simulated individuals to better reflect observed real data. We used the mean and covariance information of the individual parameters rather than the species-level parameters as the species-level distributions were built as independent and did not encode internal relationships.

The choice of Equation (4) is informed by the survey structure at Barro Colorado Island. Six observations is very little data per individual, so a unimodal function constrains the amount of wiggling that could lead to over-fitting of size dependence. Three parameters plus an initial condition is also the limit of what we can do with six data points so a more complicated function is unlikely to be appropriate. Other three-parameter hump-shaped functions would likely have similar behaviour over the observed sizes, for example a quadratic function that passes through the origin or a scaled support bump function. As these have a peak, that information could be extracted as a *g_max_* parameter even if doing so required reparameterisation. The key features there would be that the function is unimodal, hump-shaped, smooth enough to work with the Hamiltonian methods used by Stan, and has three parameters that in some fashion control the position and sharpness of the peak. As we do not see near-zero sizes or extremely large ones we cannot distinguish between the convex decay behaviour of the quadratic function, and concave decay of a bump or Canham function. We also cannot distinguish between the finite max size of quadratic/bump functions and the (technically) infinite max size of the Canham function because the trees do not get to be that big. The very asymmetric behaviour of Canham with large *k* is not distinguishable from the symmetric bump and quadratic functions from the available data as for each individual tree we do not see enough of a life history.

The choice to not include spatial dependence is practical as we wanted to provide a proof of concept with minimal covariate data. There is nothing in the hierarchical model or programming set-up that prohibits covariates, but depending on how much additional structure is included it may become prohibitively computationally expensive. The most straight-forward way to do so would be a logistic regression model with *k* covariates

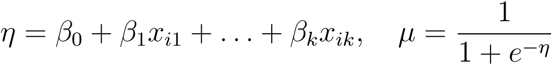

then use the output *µ* as a multiplier on *g*:

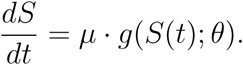

The multiplicative modification would retain the interpretation of *g_max_* as maximum potential growth rate under ideal conditions and would be interpreted as an individual-level effect as seen in Clark et al. (2007). The covariate coefficient *µ* could be interpreted as a proportion of the max potential growth, though there may be issues of identifiability in use.

From the simulated individuals we were able to demonstrate consistent error reduction for both size and growth in comparison to ‘observations’ with measurement error. The error reduction was greater for the Rüger error simulations than the N (0, 0.1) simulations as the extreme errors in Rüger simulation were rare enough to be reigned in by the longitudinal model. We suspect that the Rüger error being symmetric and centred at 0 allowed for the normal likelihood to accommodate the non-normal distribution.

Population-level parameter estimation was mixed. The hyper-parameters for *g_max_* were estimated consistently, and it’s likely that the negative bias on the mean and positive bias on the standard deviation parameters balance out. We see the same structure in the *S_max_* hyper-parameters, though the standard deviation of ln(*S_max_*) is poorly estimated overall. The hyper-parameters for *k* were difficult to estimate for the model and the positive bias on the mean indicates that individual trajectories were estimated to be flatter than the truth, which aligns with the positive bias in individual parameter estimates for *k*.

Through simulation we are able to see where individuals with near-zero growth across the observation period struggle with parameter estimation. Non-identifiability – a lack of unique parameter value that best fits the observed data (Latz, 2023) – is part of the problem for the minimum growth individual. This is not a surprise, if you have a quantity that barely changes then understanding its behaviour when undergoing change is hard. The growth parameters *S_max_* and *k* seem to be particularly sensitive to lack of observed growth. For *S_max_* we believe this is because you need to see something of either plausibly ‘high’ growth, or at least acceleration/deceleration for that individual as otherwise the MCMC must rely on the rest of the sample to constrain values. For *k*, change in growth rate is needed as it is the parameter which controls how fast that occurs which suggests that there is more to the minimum data requirements than just the number of observations.

In comparison to other growth models we have made advances in trajectory estimation and non-linear function fitting. From Herault et al. (2011), we have demonstrated considerable within-species variation in growth trajectory that cannot be observed when fitting a species average model. We are also able to get estimates of individual sizes over time which are also not accessible without fitting individual trajectories. In comparison to Clark et al. (2007) we have a size-dependent model which gives non-linear sizes over time that behave more like the observed data than a constant growth model would allow. However, we do not include a year effect as we are not including environmental covariates in order to demonstrate a minimal viable model.

There are a number of opportunities for future research building on the progress made in paper. Firstly, our growth model provides a new way to characterise growth in a species, and it is of interest to compare results across species to study ways in which growth, and intraspecific patterns in growth, vary across species. Secondly, while we were able to successfully fit our model to BCI data, such high-quality data is rare in ecology due to the difficulty of sustaining long term collections. Furthermore, simulations suggested that our method had trouble with estimation in some settings such as low growth individuals due to insufficient information, and producing a set of simulated individuals that had good convergence behaviour took some work. Deeper exploration of data requirements in order to reliably fit individual-level growth models is warranted, and simulation is a viable method for doing so.

## 6 Conclusions

We have demonstrated that a hierarchical Bayesian longitudinal model is able to improve estimates of size and growth in comparison to raw observed data, and eliminate negative growth estimates entirely through a suitable growth function and parameterisation. We have substantially improved existing methods that relied on ad hoc data exclusion through implementing a longitudinal model based on a positive growth function, and in separating individual variation from measurement error we are able to decrease the predictive error as well when compared to an approach using an average trajectory.

Our method is able to get estimates of both individual-and species-level growth function parameters. The individual level estimates are novel, and show variation within the species that has not been visible to average trajectory models. Individual parameter estimates also uncovered a way for people doing simulation studies to better represent the behaviour of a given species. The simulations and case study presented demonstrate that our proposed method is able to improve on the information available through existing population-level models and extract novel information.

## Supporting information

3D scatter plot of individual parameters

## Notes

https://github.com/Tess-LaCoil/HierarchicalLongitudinalInto

### Competing Interest Statement

The authors have declared no competing interest.

### Summary of Updates

Revisions from peer review. Clarifications to several sections with regards to things like multiple error models for simulation, and the underlying distribution structure encoded in the hierarchical method. Newly fitted models to test one-step-ahead predictive power.

