## Appendices for "Yes, they’re all individuals: Hierarchical models from repeat measurement data improve estimates of tree growth and size"

### Yes, they're all individuals: Supplementary Material

November 15, 2024

#### 1 Numerical Methods

We implemented three integration methods, as described in Butcher (2016), to investigate how they interact with the Bayesian estimation:

- Forward Euler, which has the worst performance overall but is the fastest method as it requires only one point to be computed,
- Explicit midpoint, which uses two points,
- A 4th order Runge-Kutta algorithm, which is more accurate but also computationally intensive as it has four points per step.

Both the midpoint and Euler methods are specific cases of the more general Runge-Kutta algorithm, 1st and 2nd order respectively. The Euler method is used in Iida et al. (2014).

To demonstrate the need for and impact of the different methods in size-dependent models, we simulated a single individual with growth function

$$g = \beta S, \quad \beta = 1,$$

for 5 observations with 1 unit of time between them. We added $\mathcal{N}(0, 1)$ error to each of the true sizes to simulate observations $s_j$, then fitted the longitudinal model to the data with each of the three numerical methods. The process was repeated 1000 times, where each round had different measurement error added, then all three numerical methods were fitted to the same sequence of 'observations'. Figure 1 summarises the results from this process. Panel a shows the forward projection from $S(0) = 1$ with the numerical method alone and a step size of 1. Euler and mid-point under-estimate the true solution while Runge-Kutta is difficult to distinguish from it. In Figure 1b we can see that all methods produce similar estimates of size at a particular time point when the respective integration methods are applied to data with error. To do so, the worse methods over-estimate the growth parameter $\beta$ as seen in Figure 1c, where only RK4 is able to accurately estimate the true value of 1. By over-estimating $\beta$, Euler and midpoint are able to haul the forward projected sizes up to meet the required solution. As we are interested in estimating parameters, we must use the most accurate method and progress with RK4.

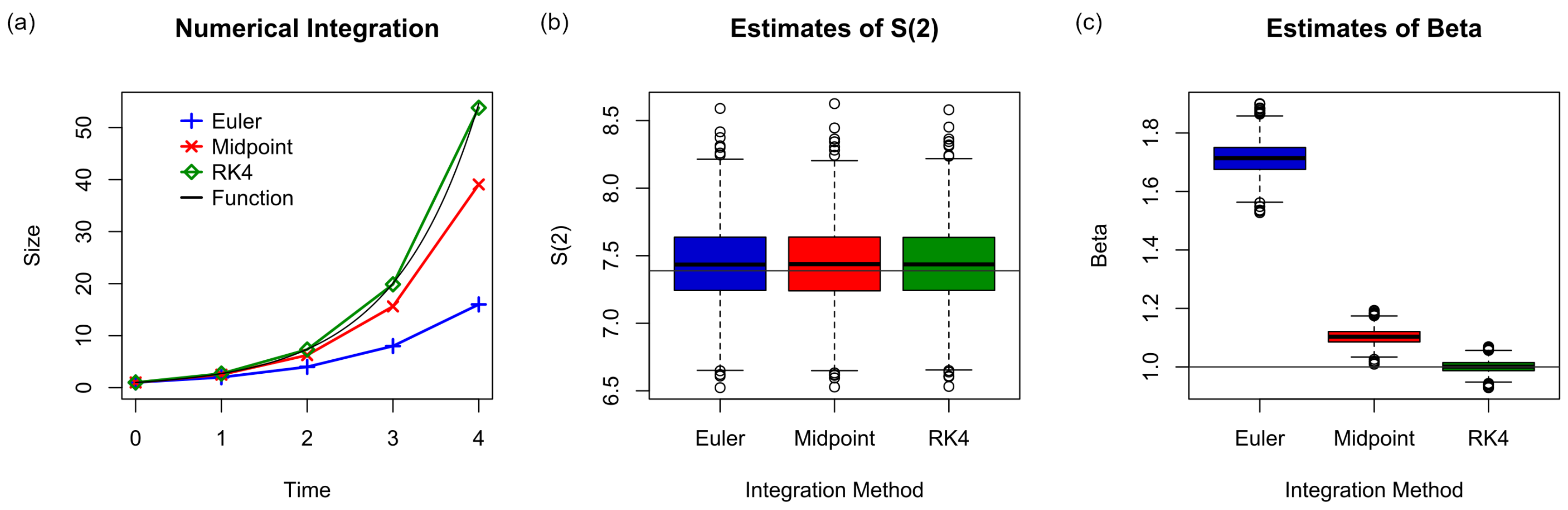


Figure 1: Demonstration of the role that numerical methods play in accurate parameter estimation. **(a)** Demonstration of the different numerical integration methods and the varying fit to the analytic solution without the Bayesian model. We provide $\beta = 1$ to each. **(b)** Box plots of estimated values for $S(2)$ arising from 1000 simulations of measurement error, each of which was fit by all methods. The solid like indicates the true size. **(c)** Box plots of $\beta$ estimates from 1000 simulations, fit by each method. The solid line indicates the true value of 1. Only RK4 gives reliable estimates, as the other two have strong positive bias.

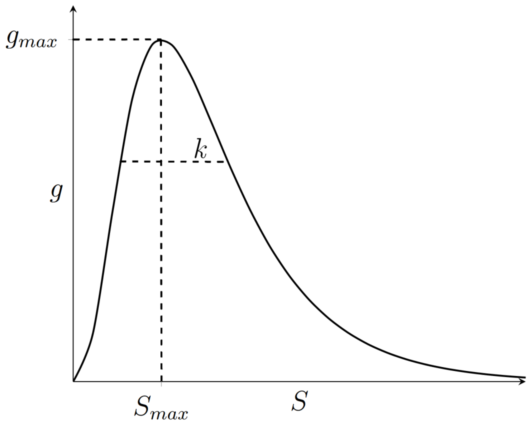


Figure 2: Impact of parameters for the Canham model, where $g_{max}$ determines the maximum growth rate, $S_{max}$ controls the size when that peak occurs, and $k$ controls how rapidly growth declines away from the peak.

### 2 Canham Details

What we describe as the Canham growth function is fully parameterised by three paramters:

- $g_{max}$ the maximum growth rate,
- $S_{max}$ the size at which that peak occurs,
- $k$ which controls the peakedness of the curve by both the decay in growth after the peak, and how rapid the acceleration is to the peak.

Of these, $k$ is the most difficult to conceptualise, we think of it as a spread parameter where larger values mean a gentler peak, small values lead to spikes in growth.

### 3 BCI Diagnostics

At the level of individual size trajectories all the estimates were reasonable to the underlying data based on visual comparison of estimated and observed sizes over time.

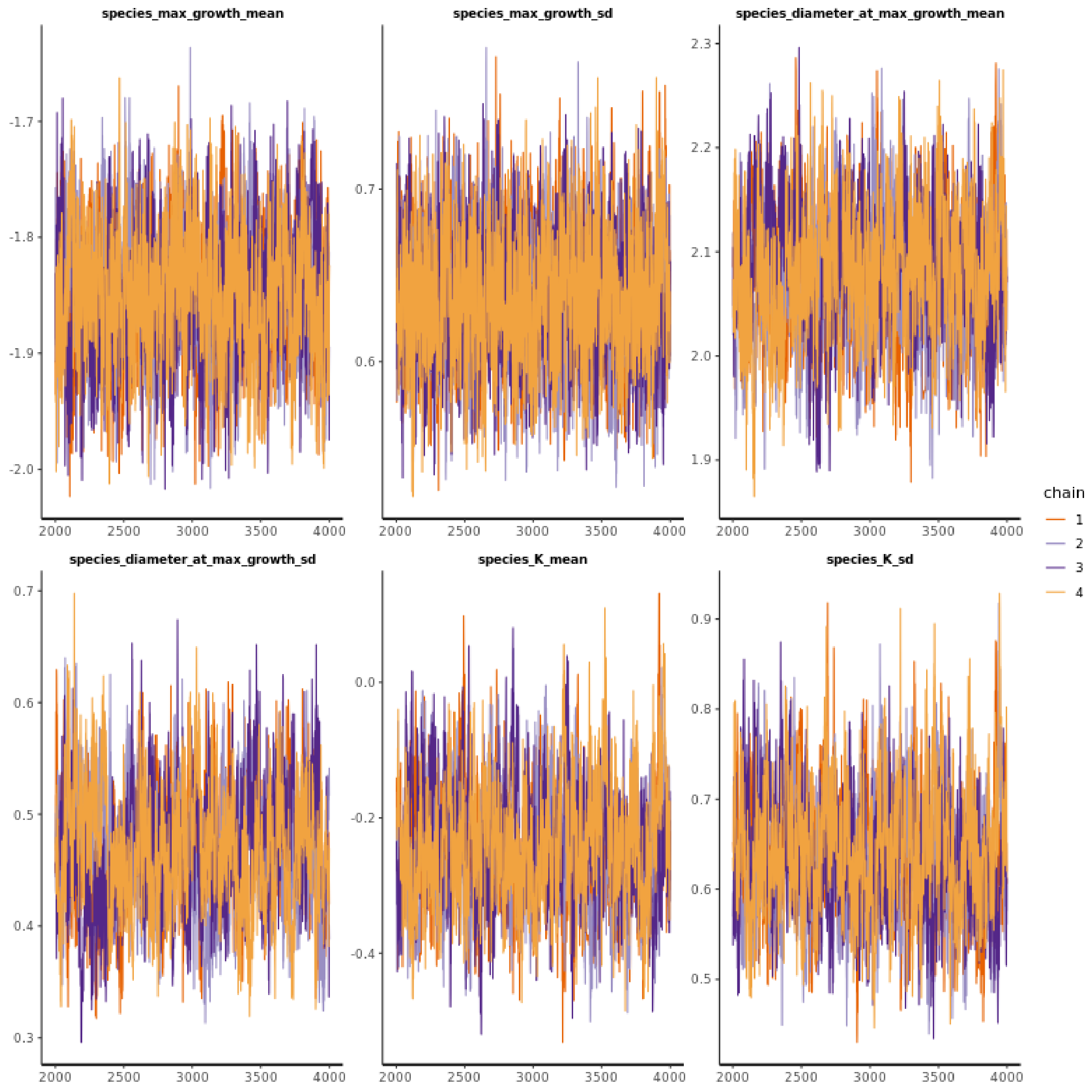


Figure 3: BCI demonstration data with Canham model. Traceplot of chains for species-level hyper-parameters.

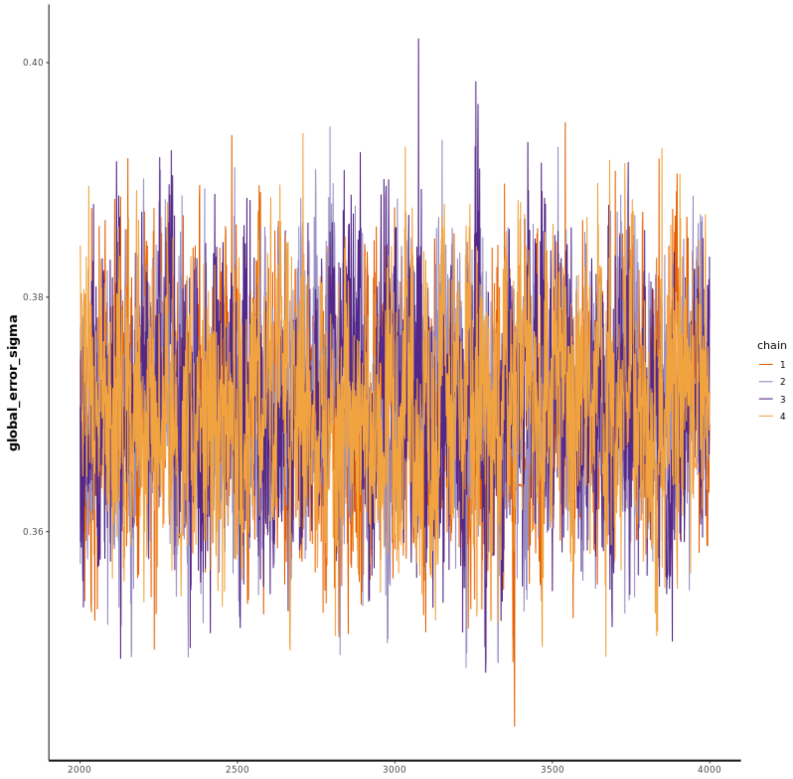


Figure 4: BCI demonstration data with Canham model. Traceplot of chains for global error parameter.

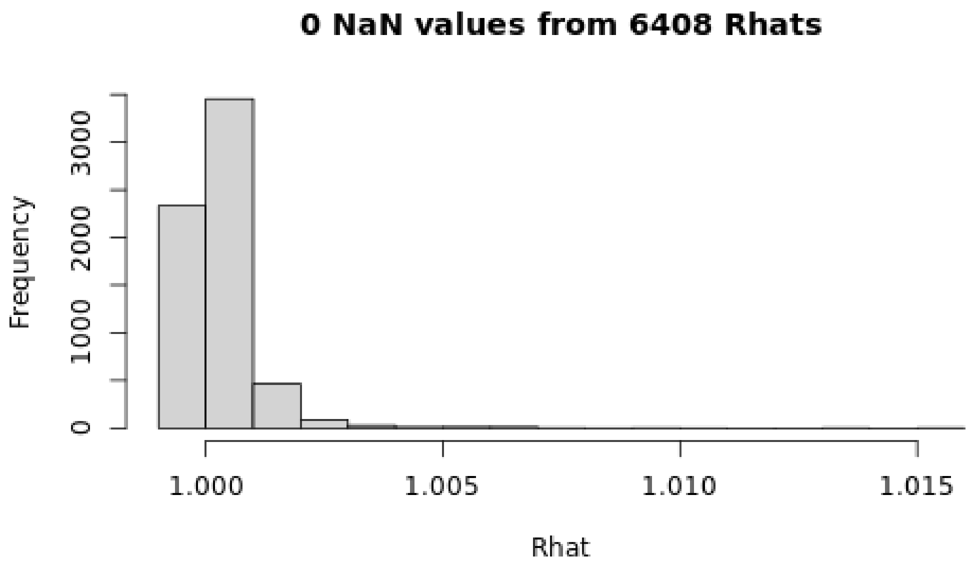


Figure 5: Histogram of $\hat{R}$ for all chains of BCI demonstration model with Canham model.

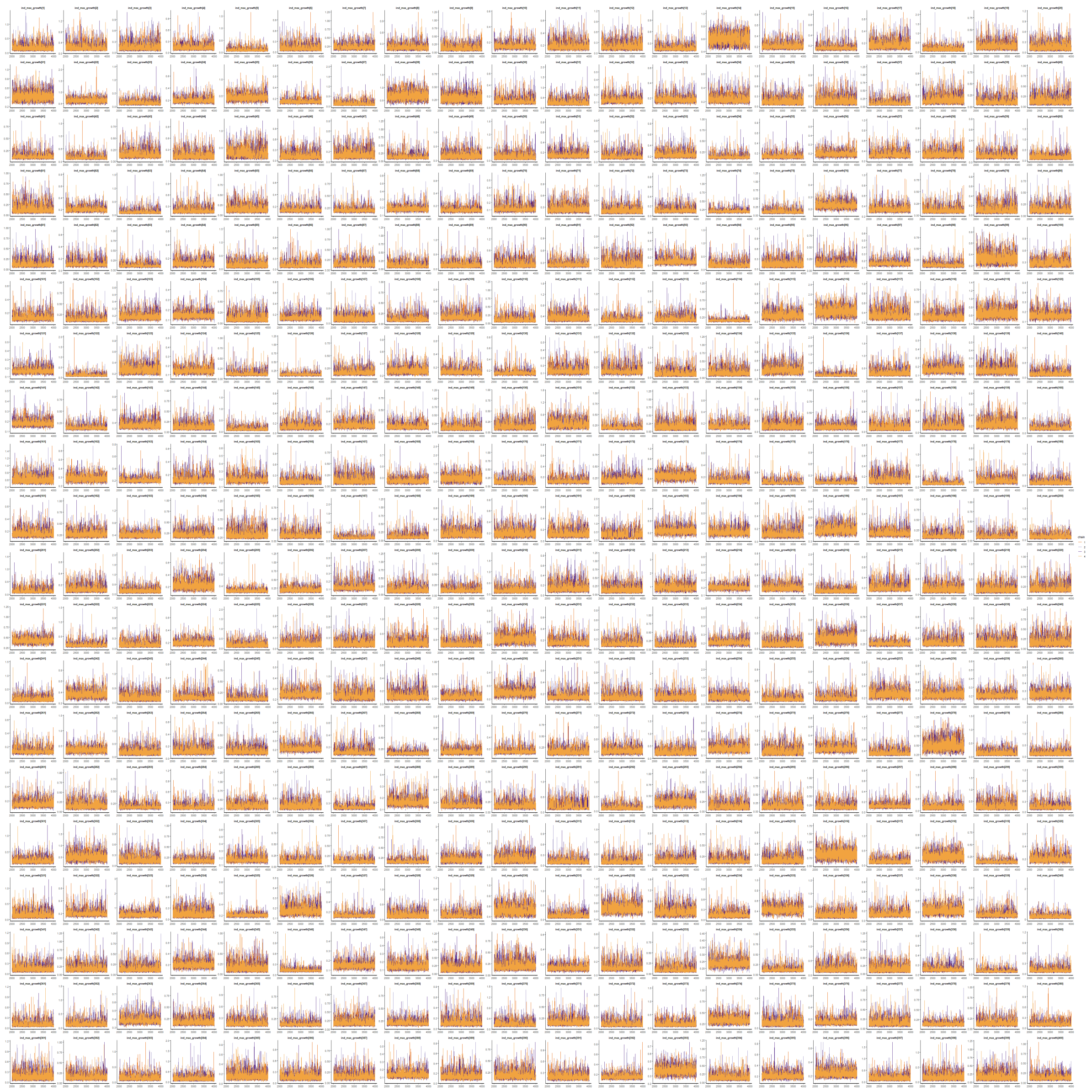

Figure 6: Diagnostic plots for individual-level $g_{max}$ parameters.

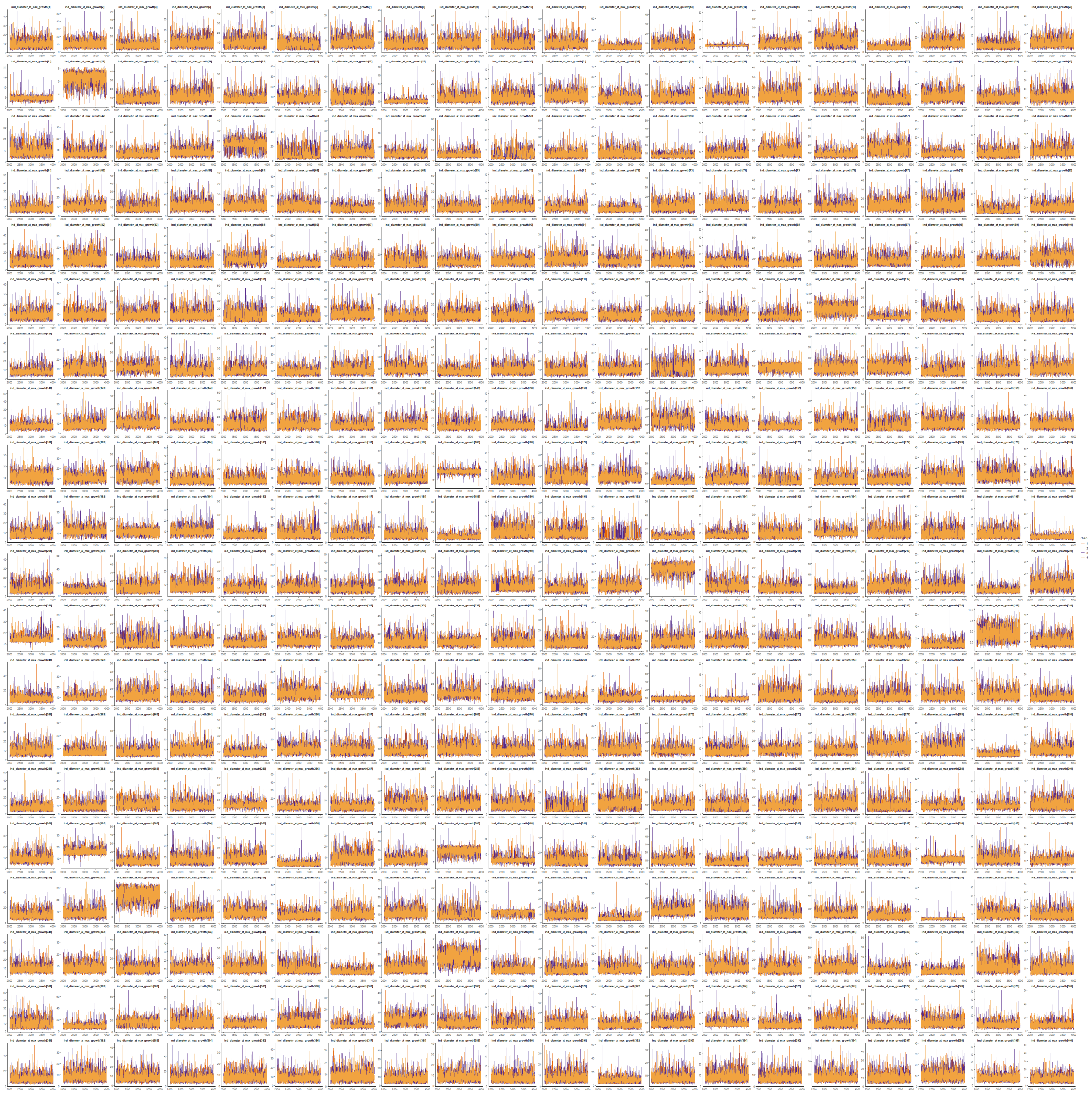

Figure 7: Diagnostic plots for individual-level $S_{max}$ parameters.

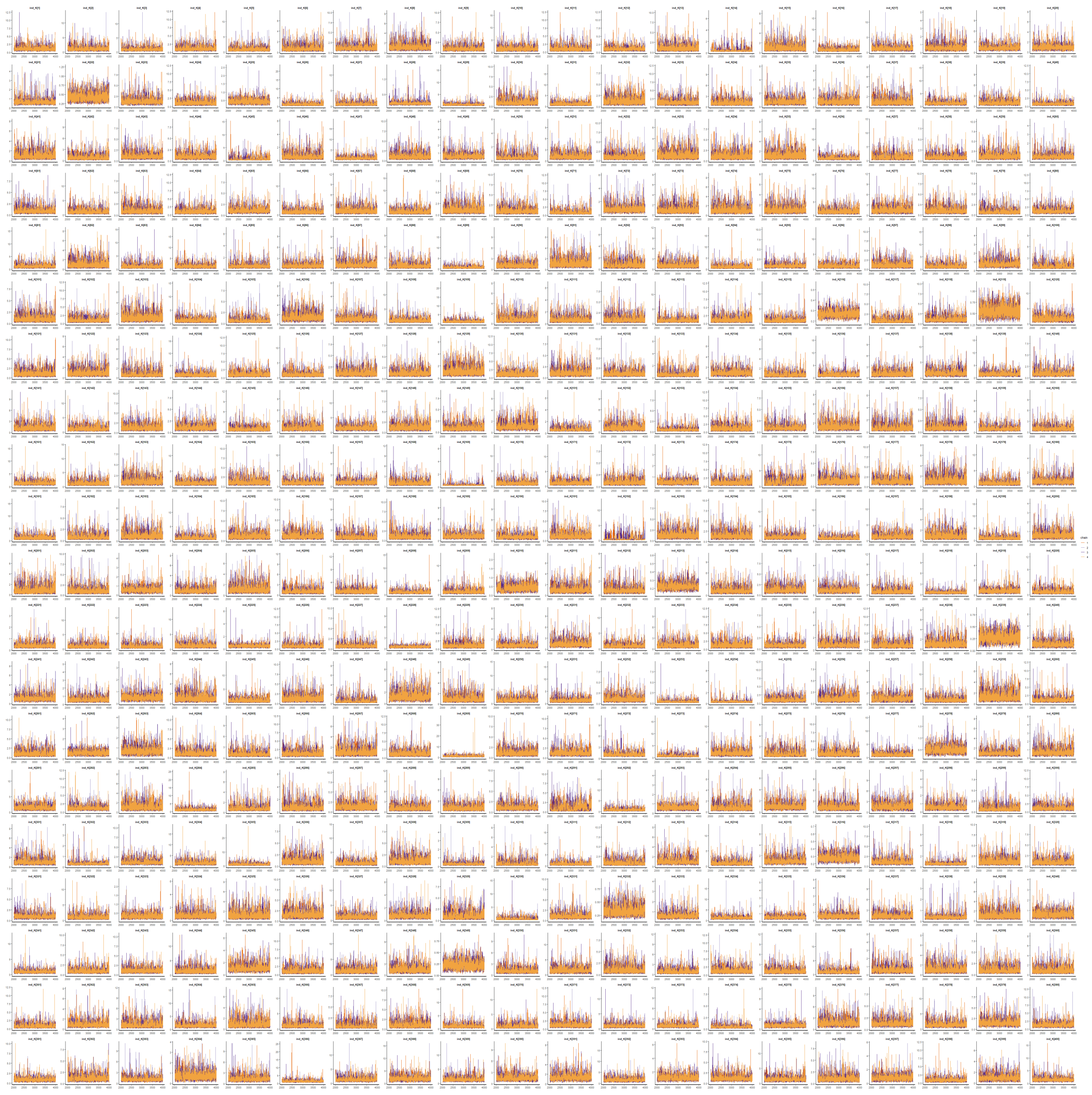

Figure 8: Diagnostic plots for individual-level $k$ parameters.

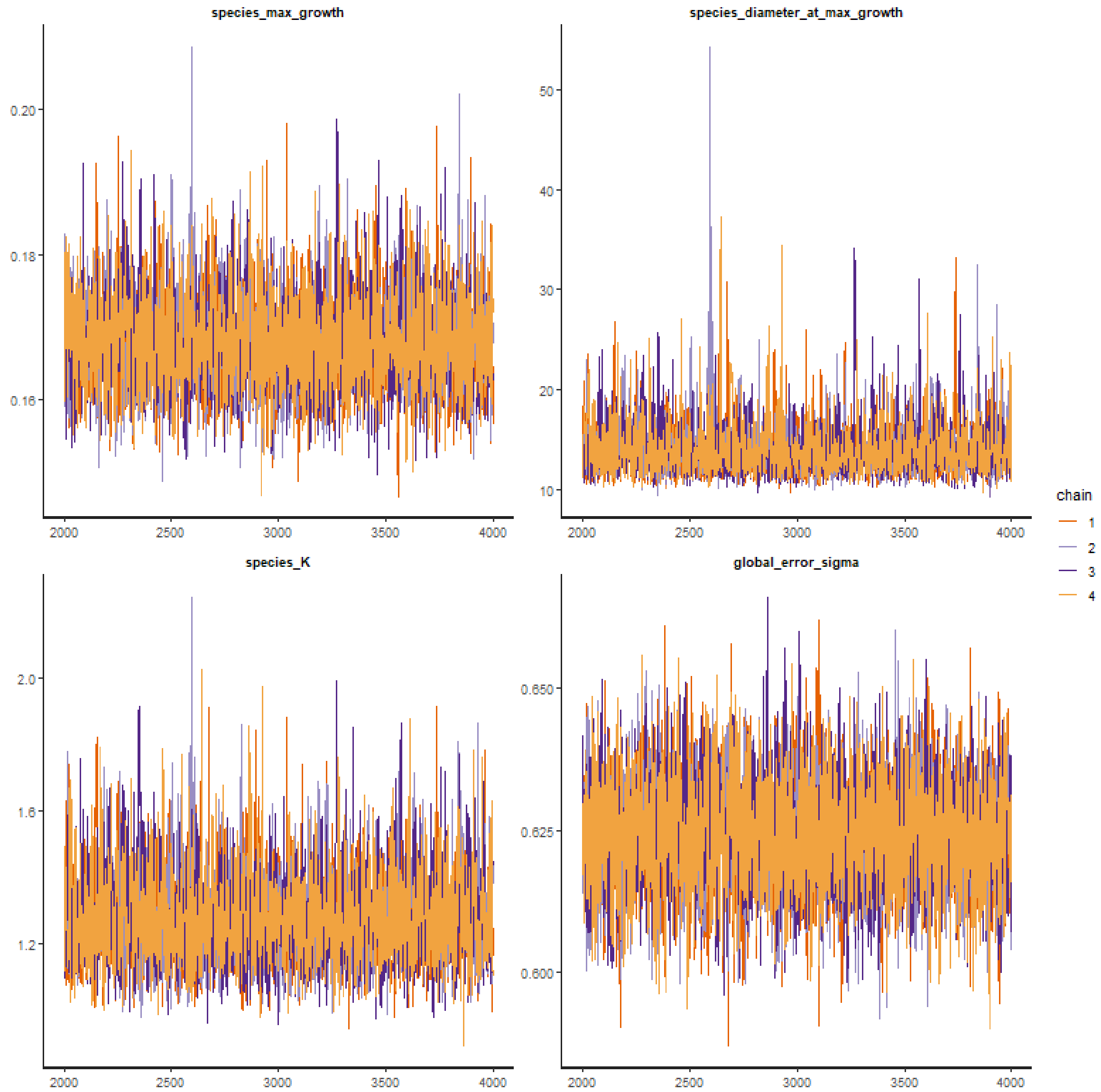


Figure 9: Diagnostic plot for species-level model fit to pairwise differences.

### 4 BCI Additional Results

We want to look at the individual parameters in greater detail. Figure 10 gives distribution information in one and two dimensions. See the supplementary material for a 3-dimensional scatter plot that is useful for examining the population structure. From Panels (a), (b), and (c) we can see that there is evidence of relationships between parameters, and regions of the parameter space which are not observed.

Small $S_{max}$ and small $g_{max}$, which would correspond to peak growth at a small size, tend not to occur together, and there is preliminary evidence of a negative relationship between the two parameters. There seems to be a positive relationship between $k$ and $g_{max}$ for trees with lower $g_{max}$ values, but those with the highest peaks have a steep drop-off in $k$, which corresponds to a sharper peak in growth.

In 10(c) we have the clearest indication of a relationship, with what appears to be a negative straight line relationship in the log-transformed scale. High $S_{max}$ and lower $k$ would produce a peak late in size with the Canham function, however the $k$ values observed for the bulk of the population are not so small that we see a very narrow peak, which aligns with the fitted growth curves we see in Figure 3 from the main paper.

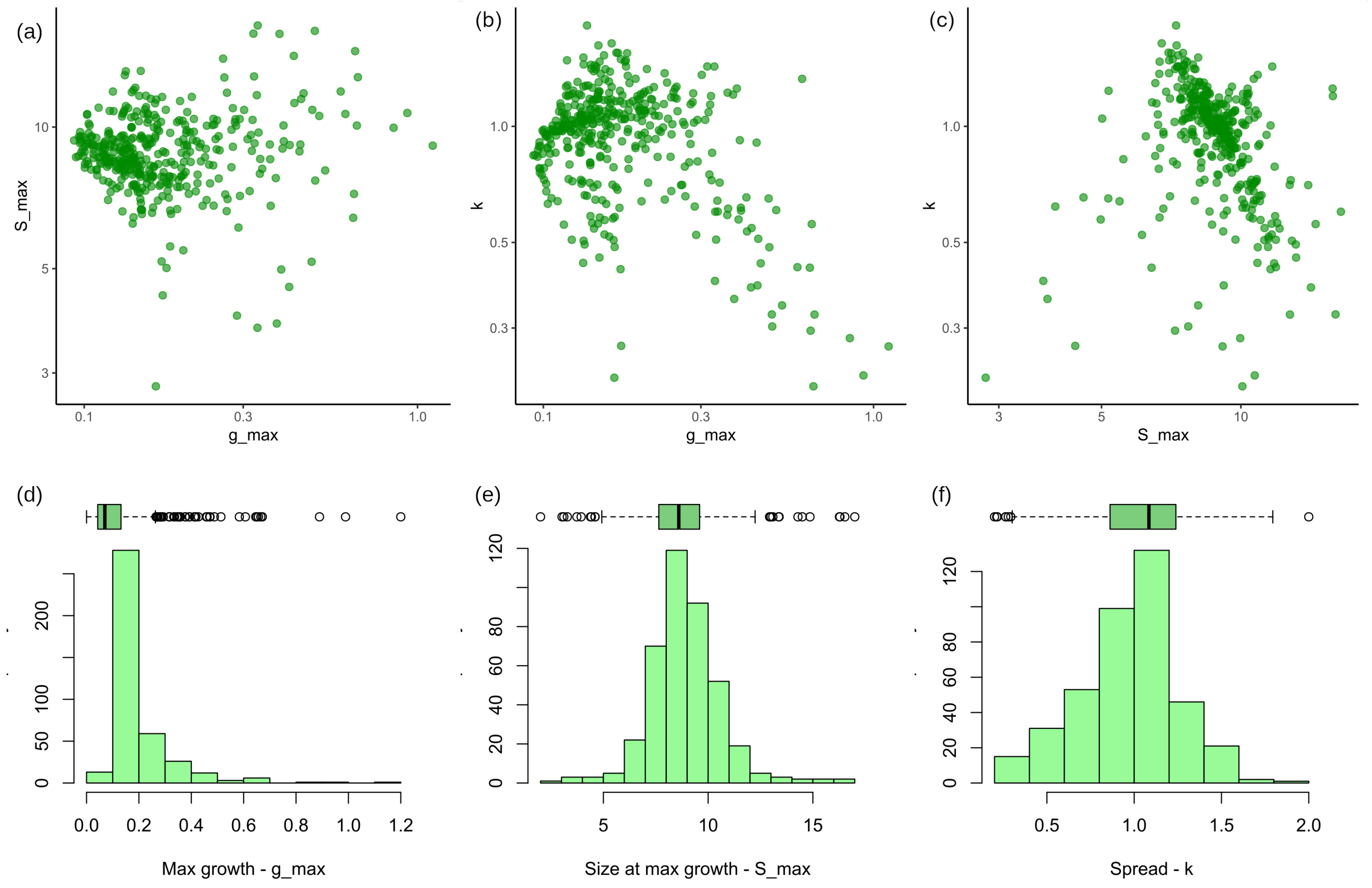


Figure 10: Scatterplots and histograms of the individual parameter estimates from *G. recondita* showing possible relationships between the individual parameter estimated values.

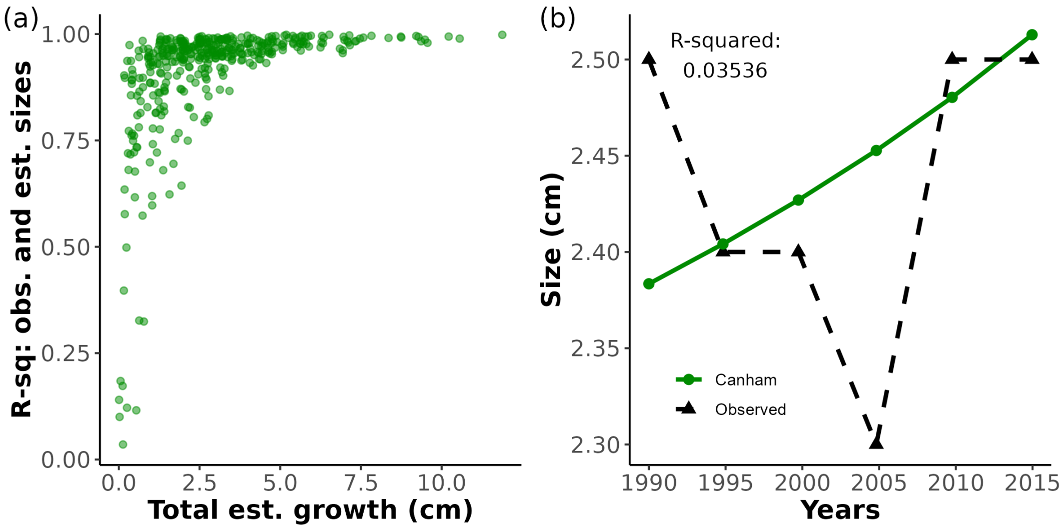


Figure 11: **(a)** The distribution of $R^2$ statistics for each individual fit shows that across the sample the relationship between fitted and observed sizes is very strong. Individuals with low overall growth are the only ones that demonstrate low $R^2$. **(b)** is the individual with smallest $R^2$, and demonstrates the negative increments and low overall growth typical of these trees.

#### 4.1 Results for $R^2$ distribution

Figure 11(a) shows the relationship between estimated total growth and $R^2$ of estimated and observed sizes. Of 400 trees, 33 had $R^2 < 0.75$, all of which had estimated total growth less than 2.2cm. Seven trees had $R^2 < 0.25$.

### 5 One-step-ahead estimation

To test the out-of-sample fit capacity of our model we two growth functions to the first five *G. recondita* observations and used the fitted growth function for each individual to estimate the 6th size:

1. The hierarchical Canham model fit to 6 observations that directly estimates $\hat{S}_{i,6}$,

2. A hierarchical Canham model fit to the first 5 observations that extrapolates $\hat{S}_{i,6}$ from $\hat{S}_{i,5}$ using the individual-level growth parameters,

3. A hierarchical model with constant growth rate fit to the first 5 observations that extrapolates $\hat{S}_{i,6}$ from $\hat{S}_{i,5}$ using the individual-level growth parameter.

The constant growth rate model is directly analogous to a linear mixed-effects model for sizes over time with individual term. We chose this as our closest equivalent model for predicting sizes as the species-average growth model of Herault et al. (2011) is not used for size prediction, but hierarchical models with linear sizes over time are, for example Zhao et al. (2013) which is looking at size-dependent growth in fir.

Figure 12 shows that of the two models fit to the first five observations, the hierarchical Canham model fit the best with an RMSE of 0.69 compared to 0.787. The constant model also had a more left-skewed distribution of errors, indicating more under-estimation of observed final sizes which is likely related to the decline of growth at large sizes.

#### 5.1 Diagnostics for one-step-ahead models

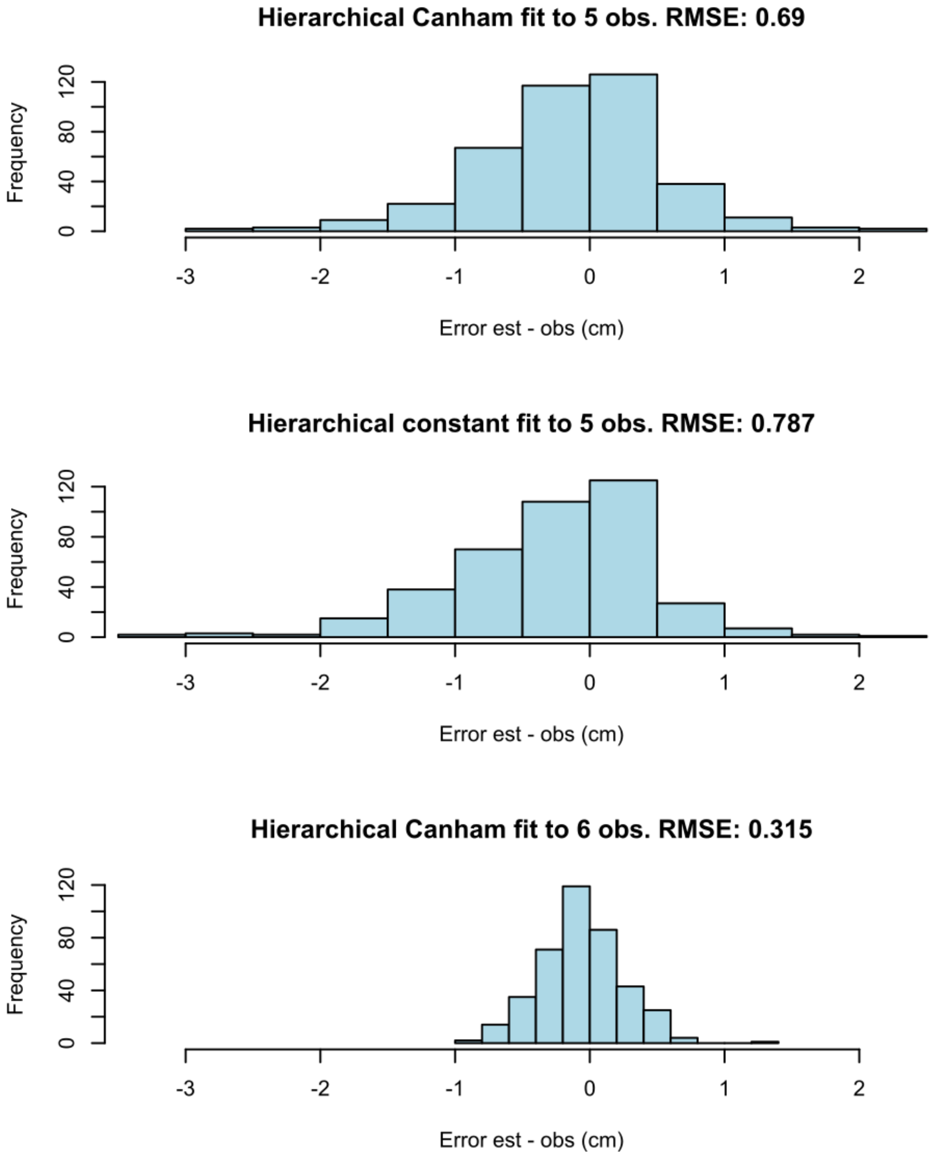


Figure 12: Comparison of the difference between estimated and observed final sizes for *G. recondita* based on different models. We see that the direct estimation of $\hat{S}_{i,6}$ in the hierarchical Canham model fit to 6 observation had the lowest RMSE, followed by the hierarchical Canham model fit to the first 5 observations. The alternate method – a hierarchical model fitting linear sizes over time with an individual effect – performed less well than either of the non-linear growth functions.

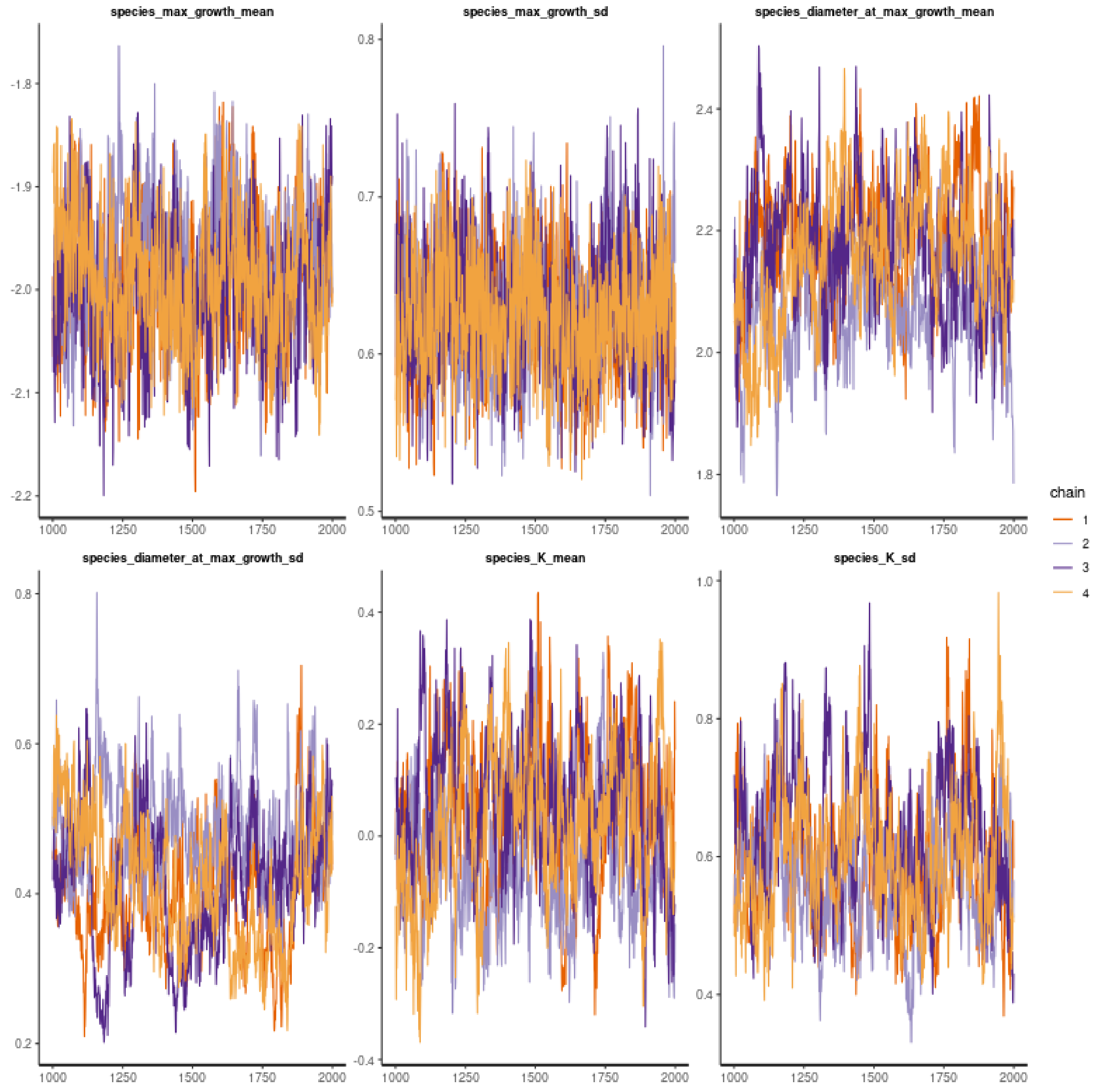


Figure 13: BCI demonstration data with Canham model fit to 5 observations. Traceplot of chains for species-level hyper-parameters.

### 6 Simulation Sample Information

To reduce factors varying across the simulations we first built a set of 50 individuals whose true sizes would form the underlying data for all simulations. As there can be problems with badly behaved parameter value combinations, we ran small models on batches of 10 individuals at a

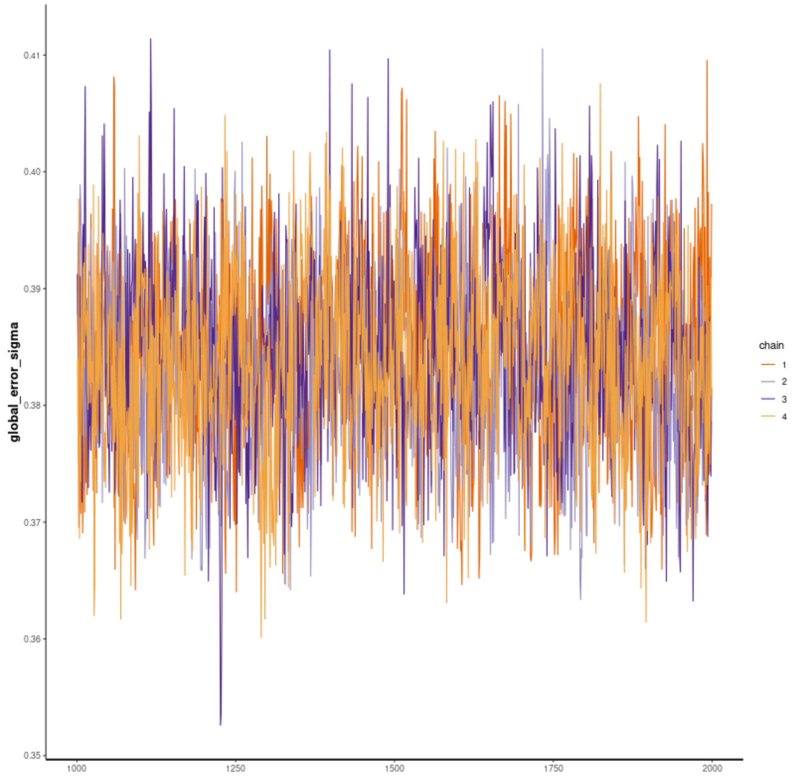


Figure 14: BCI demonstration data with Canham model fit to 5 observations. Traceplot of chains for global error parameter.

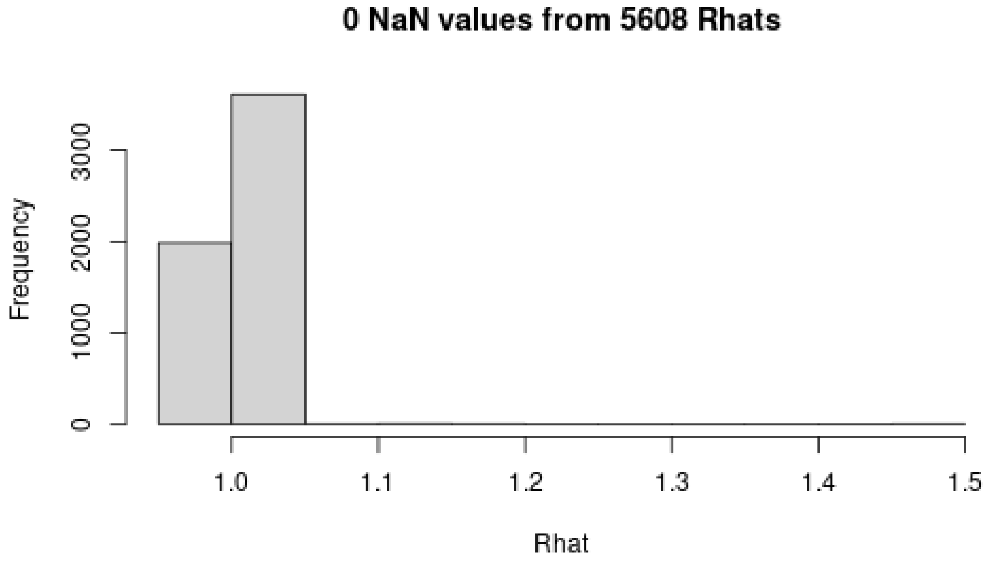


Figure 15: Histogram of $\hat{R}$ for all chains of BCI demonstration model with Canham model fit to 5 observations.

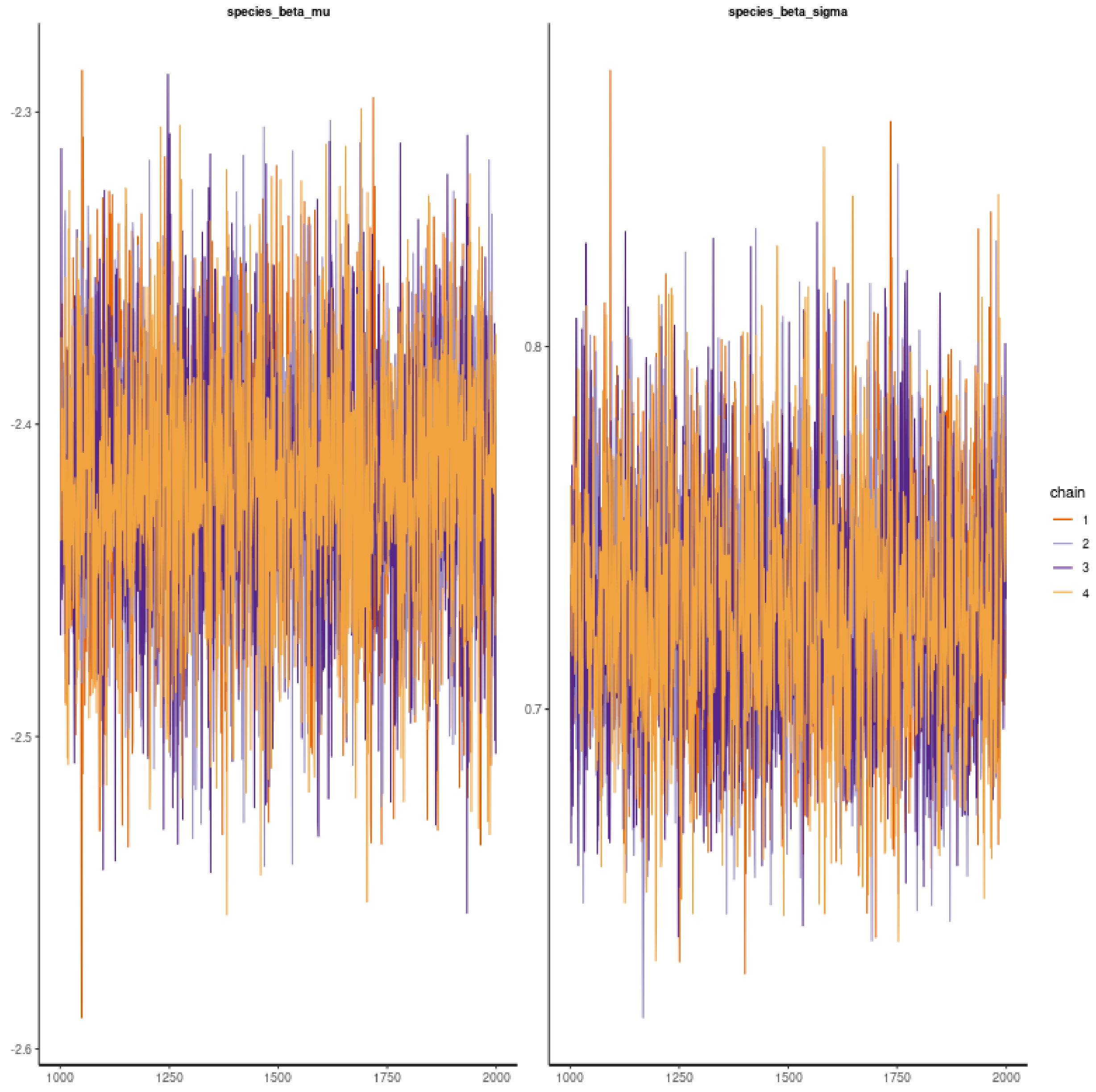


Figure 16: BCI demonstration data with constant model fit to 5 observations. Traceplot of chains for species-level hyper-parameters.

time. We excluded any that produced sampling chains with estimates orders of magnitude beyond the true parameter values as we did not observe this behaviour in the *G. recondita* fit, as can be seen in Appendix 3.

Figure 19 gives a visual comparison of the 50 simulated individuals and the *G. recondita* model, and

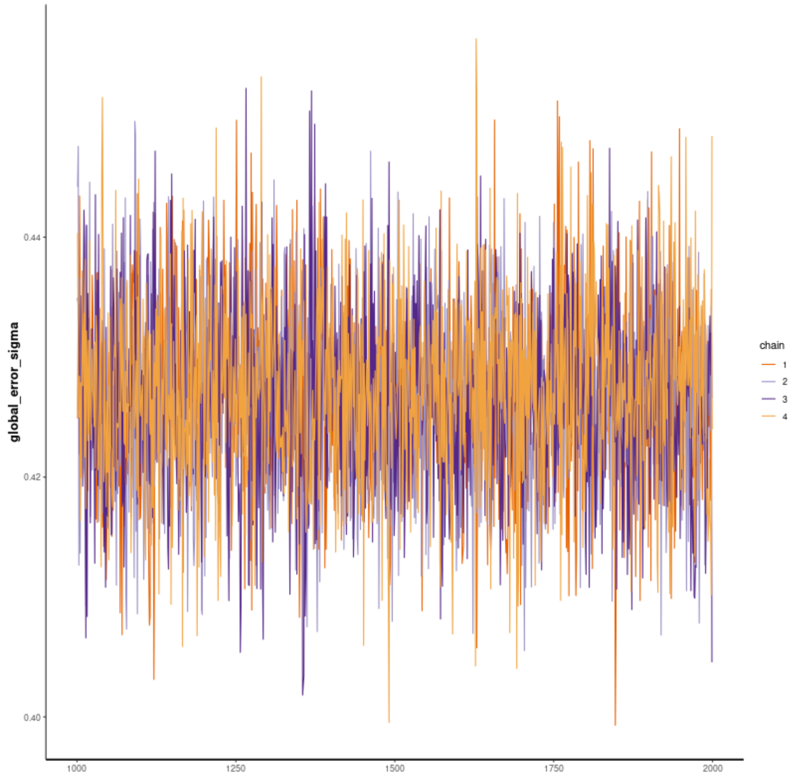


Figure 17: BCI demonstration data with constant model fit to 5 observations. Traceplot of chains for global error parameter.

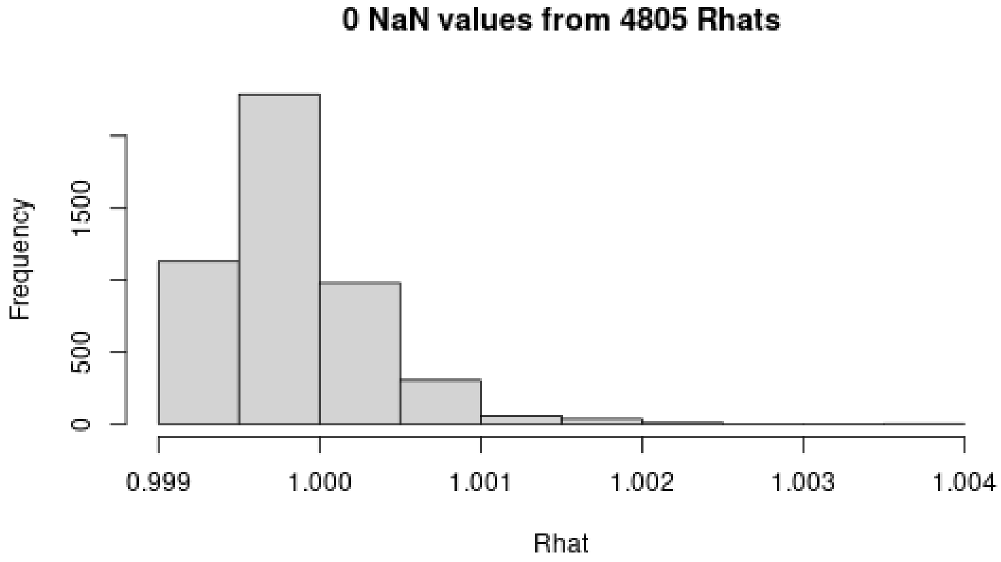


Figure 18: Histogram of $\hat{R}$ for all chains of BCI demonstration model with constant model fit to 5 observations.

Table 1 shows how the means and covariances compare. We used the individual-level distribution rather than the species-level parameters to capture the covariance structure as we have evidence that parameters are not independent.

From Figure 19 we can see that the visible parts of individual growth functions do not include much of a drop-off at larger sizes, which plausibly makes it harder to estimate larger $k$ values.

Table 1: Population and sample variances, covariances, and means for log-transformed individual parameters in simulated 50 individual Canham sample. We compare model estimates to the sample values as the 'true' parameters rather the population values for the distribution to avoid biasing results.

| | | |
|---|---|---|
| $\sigma^2_{\ln(g_{max})} = 0.199$ | $\mu_{\ln(g_{max})} = -1.769$ | $\mu_{\ln(S_{max})} = 2.177$ |
| $\sigma_{\ln(g_{max})}\sigma_{\ln(S_{max})} = 0.00457$ | $\sigma^2_{\ln(S_{max})} = 0.0368$ | $\mu_{\ln(k)} = -0.0597$ |
| $\sigma_{\ln(g_{max})}\sigma_{\ln(k)} = -0.0637$ | $\sigma_{\ln(S_{max})}\sigma_{\ln(k)} = -0.0134$ | $\sigma^2_{\ln(k)} = 0.128$ |
| $s^2_{\ln(g_{max})} = 0.174$ | $\bar{x}_{\ln(g_{max})} = -1.323$ | $\bar{x}_{\ln(S_{max})} = 2.170$ |
| $s_{\ln(g_{max})}s_{\ln(S_{max})} = 0.0116$ | $s^2_{\ln(S_{max})} = 0.0236$ | $\bar{x}_{\ln(k)} = -0.238$ |
| $s_{\ln(g_{max})}s_{\ln(k)} = -0.081$ | $s_{\ln(S_{max})}s_{\ln(k)} = -0.0037$ | $s^2_{\ln(k)} = 0.153$ |

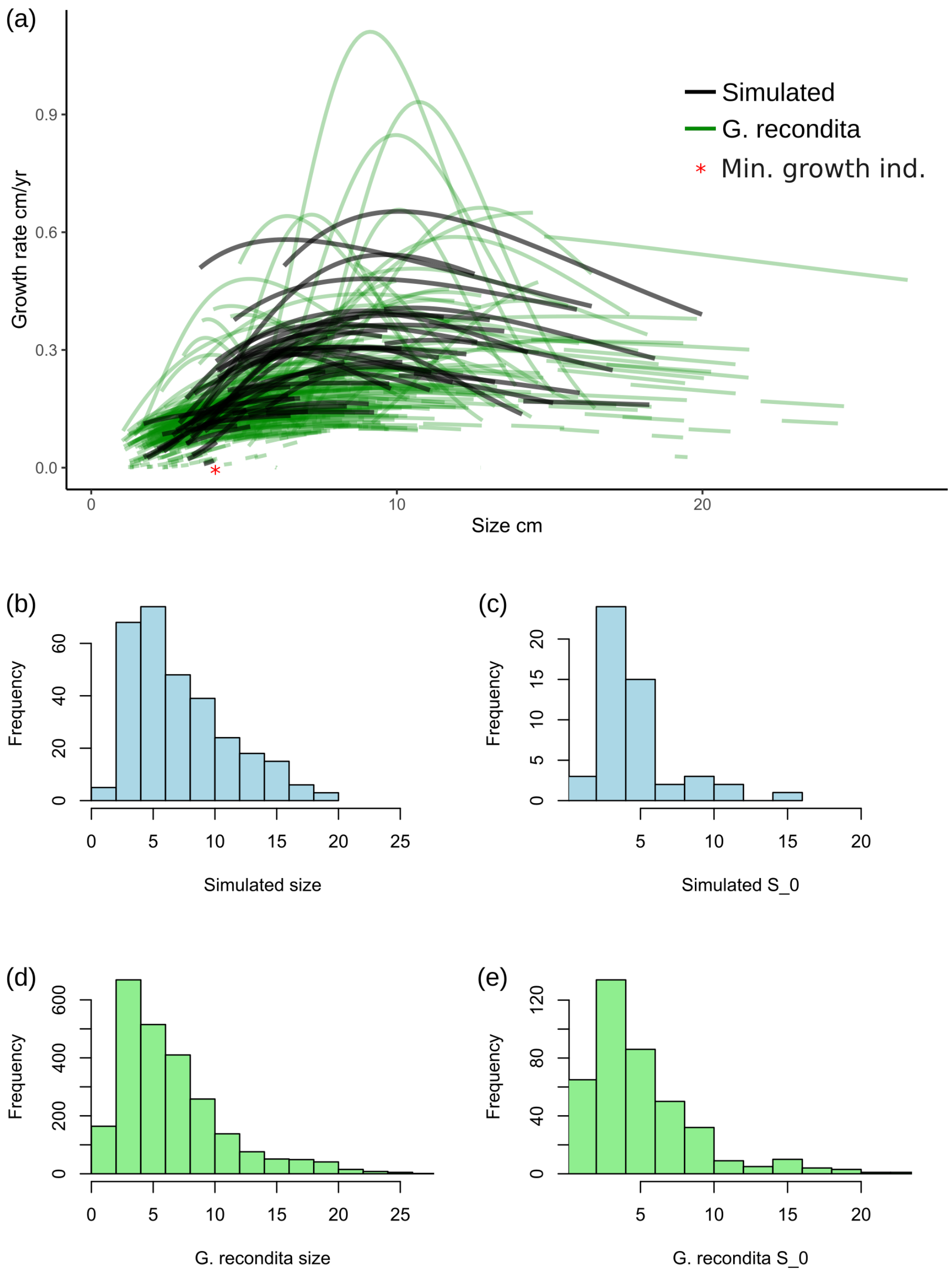


Figure 19: Comparison of simulated data without error to *G. recondita* fitted model showing that simulations align with real data. **(a)** shows that the simulated growth functions are reasonable given the observed data, and indicates the growth curve for the simulated individual with the least growth: No. 35 with 0.33 cm total growth. **(b)** and **(d)** show that the simulated sizes are a reasonable match to those observed, though there are fewer large values in the simulated data. **(c)** and **(e)** show that the initial sizes are comparable, though there are fewer large values in the simulated data.

### 7 Simulation Additional Results

Table 2: RMSE across 1000 simulation for estimates of growth and size comparing model fit to observed size (with measurement error) and pairwise difference in sizes. The Bayesian method shows considerable improvement in estimates for growth, and more modest improvement in estimates for size.

| | | **Mean RMSE** | | **Mean RMSE** |
|---|---|---|---|---|
| **Error Type** | **Measure** | Without model | Bayesian | **% Reduction** |
| $\mathcal{N}(0, \sigma_e = 0.1\text{cm})$ | Size | 1.73 | 1.22 | 29.4% |
| | Growth | 2.24 | 0.87 | 61.1% |
| Ruger Eqn. | Size | 2.10 | 1.46 | 30.3% |
| | Growth | 2.70 | 1.01 | 62.7% |

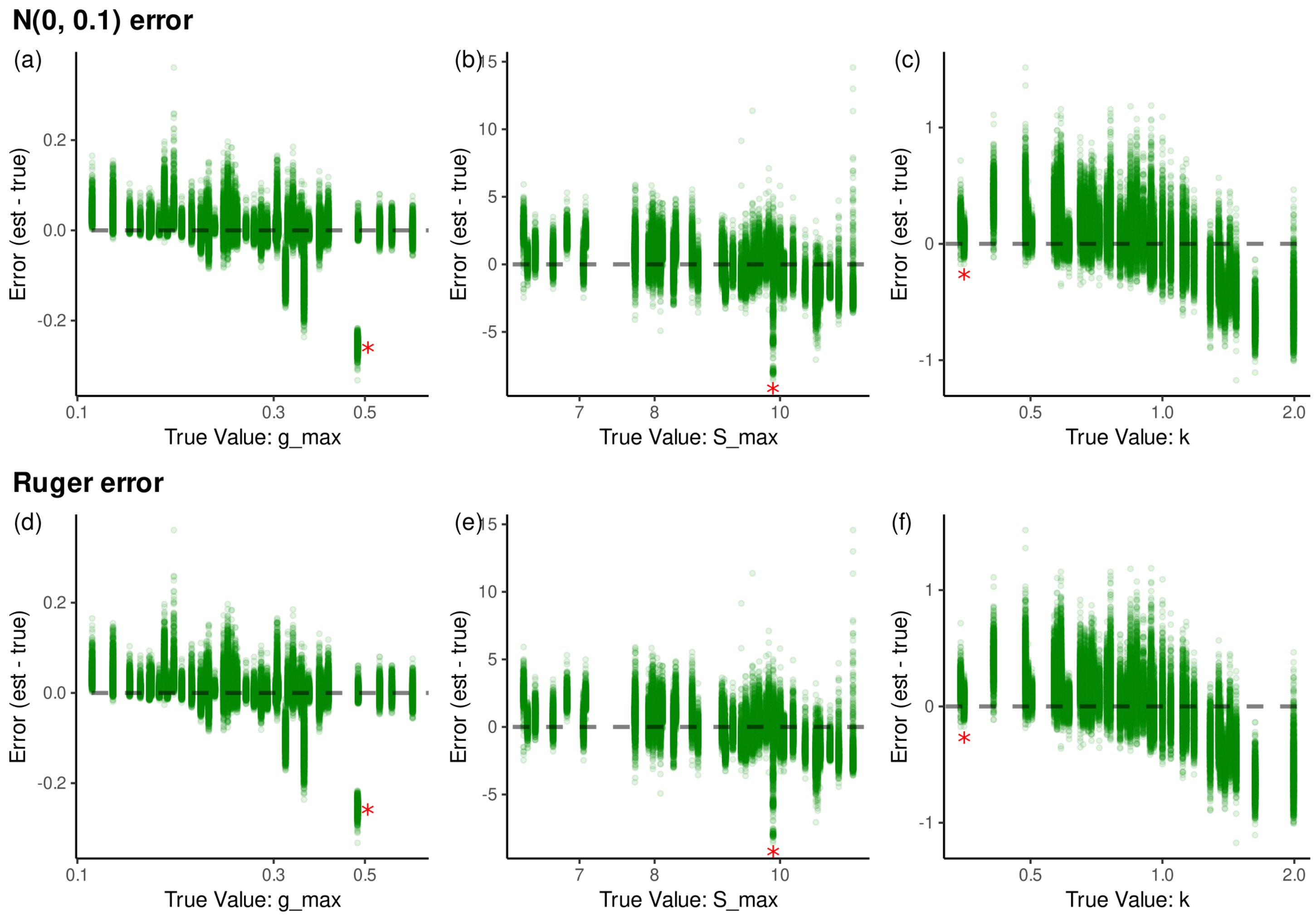


Figure 20: Scatter plots of individual parameter estimate errors with $\mathcal{N}(0, 0.1)$ and Rüger model (Rüger et al., 2011) as measurement error processes. The dashed line indicates 0 error where the estimate matches the true value. The red asterisk points to the estimates for individual with smallest total growth, which in (a) is the lower of the two estimate clusters at that value only.

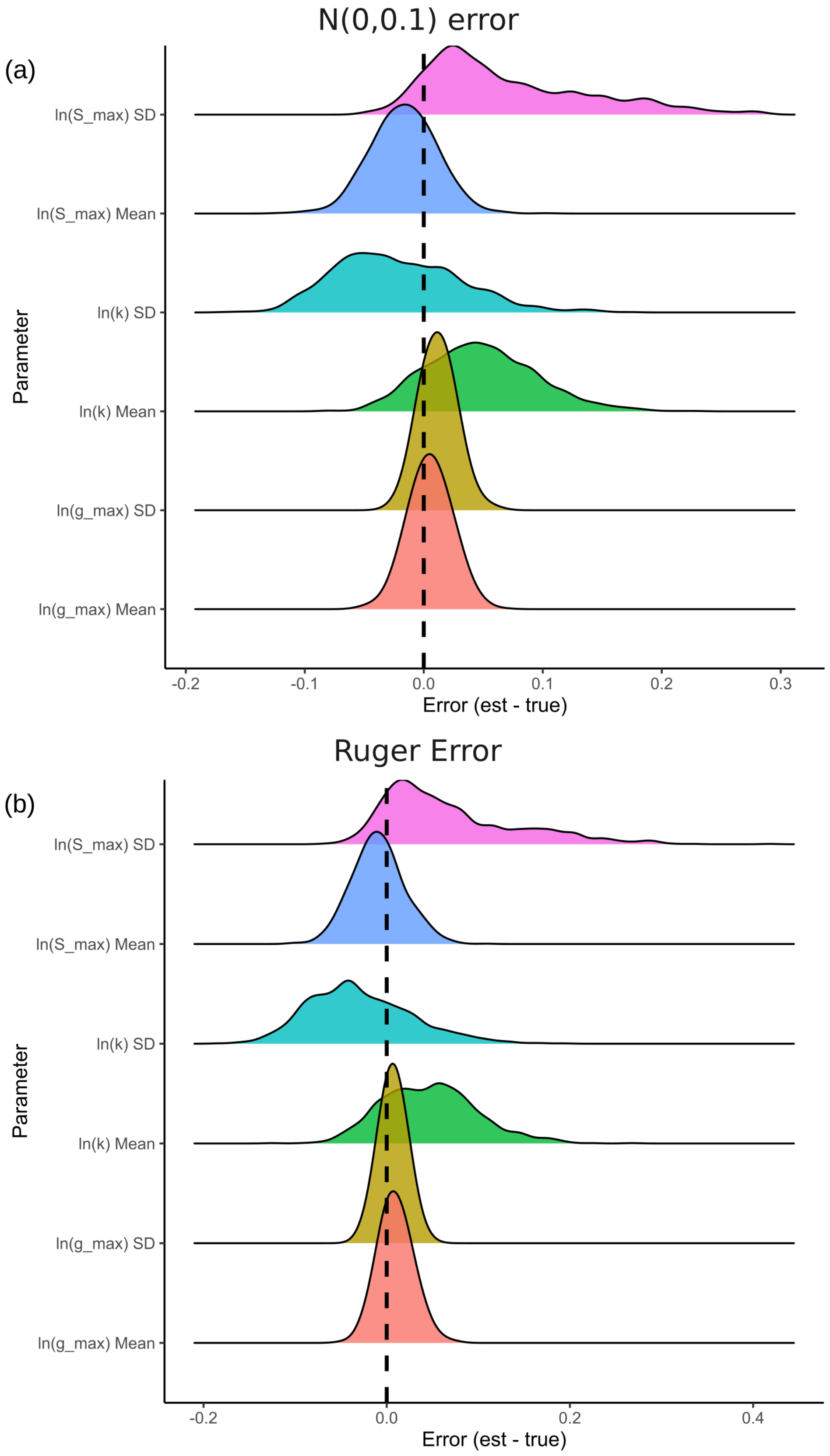


Figure 21: Density plots of errors in species-level parameter estimation. Dashed line indicates 0 error.

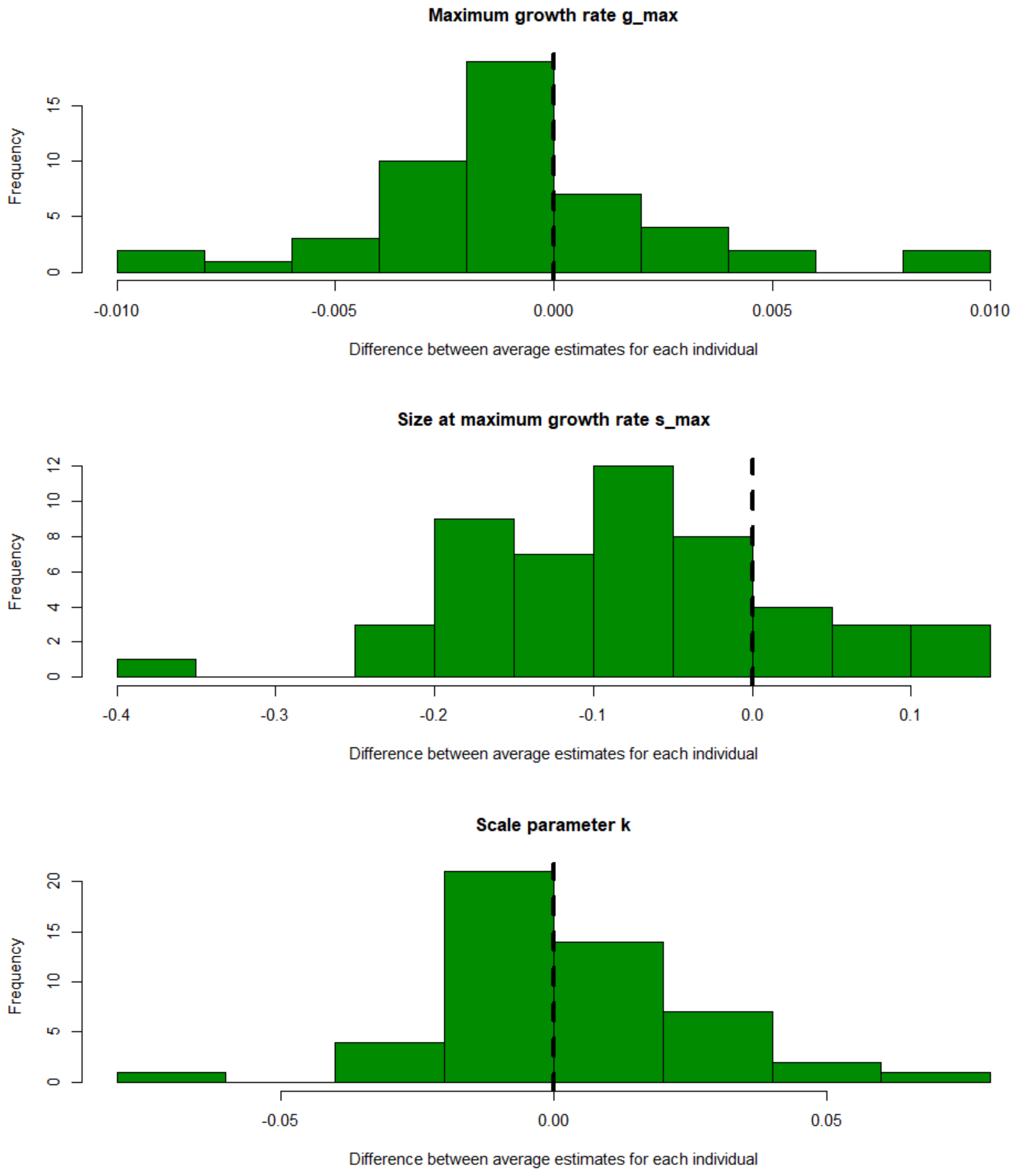


Figure 22: Histogram of the difference of means of individual parameter estimates across two error models.

#### 7.1 Simulation 3: Impact of sample size on population parameter estimates

In order to investigate the impact of sample size on population parameter estimation we simulated from a constant growth model

$$g = \beta, \quad \beta_i \sim \ln\mathcal{N}(\mu_\beta = -1, \sigma_\beta = 1)$$

across four sample sizes ($n = 10,\ 100,\ 1000,\ 10,000$). We used an additive error model

$$s_{ij} = S_{ij} + \mathcal{N}(0, \sigma_e), \quad \sigma_e = 0.2\text{cm}$$

with the standard deviation taken from the Rüger et al. (2011) size-dependent error with size of 30 cm. Constant growth was chosen as a single individual growth parameter allows for fast model runs. Each individual had five observations, with a 5 year observation interval.

Table 3 gives a summary for 1000 simulations at each sample size. The coverage of 95% posterior credible intervals was good across the sample sizes, and the mean CI width decreased by a factor of $1/\sqrt{n}$ as the sample size increased.

Table 3: Parameter estimates for constant growth model across different sample sizes and the computational time required for each. Run time includes the time to simulate data and load the stan model.

| **Parameter** | **Sample Size** | **Mean Estimate** | **95% CI Coverage** | **Mean CI Width** | **Mean Run Time** |
|---|---|---|---|---|---|
| $\mu_\beta = -1$ | 10 | -0.886 | 93.9% | 1.29 | 1.4 mins |
| | 100 | -0.991 | 94.5% | 0.393 | 2 mins |
| | 1,000 | -0.998 | 94.3% | 0.124 | 6 mins |
| | 10,000 | -1.000 | 94.9% | 0.0392 | 2 hrs 20 mins |
| $\sigma_\beta = 1$ | 10 | 1.063 | 95.9% | 1.055 | – |
| | 100 | 1.005 | 94% | 0.283 | – |
| | 1,000 | 1.002 | 94.9% | 0.0881 | – |
| | 10,000 | 1.001 | 95% | 0.028 | – |
